# Lactate is a major energy substrate for cortical neurons and enhances their firing activity

**DOI:** 10.1101/2021.05.17.444414

**Authors:** Anastassios Karagiannis, Thierry Gallopin, Alexandre Lacroix, Fabrice Plaisier, Juliette Piquet, Hélène Geoffroy, Régine Hepp, Jérémie Naudé, Benjamin Le Gac, Richard Egger, Bertrand Lambolez, Dongdong Li, Jean Rossier, Jochen F. Staiger, Hiromi Imamura, Susumu Seino, Jochen Roeper, Bruno Cauli

## Abstract

Glucose is the mandatory fuel for the brain, yet the relative contribution of glucose and lactate for neuronal energy metabolism is unclear. We found that increased lactate, but not glucose concentration, enhances the spiking activity of neurons of the cerebral cortex. Enhanced spiking was dependent on ATP-sensitive potassium (K_ATP_) channels formed with Kir6.2 and SUR1 subunits, which we show are functionally expressed in most neocortical neuronal types. We also demonstrate the ability of cortical neurons to take-up and metabolize lactate. We further reveal that ATP is produced by cortical neurons largely via oxidative phosphorylation and only modestly by glycolysis. Our data demonstrate that in active neurons, lactate is preferred to glucose as an energy substrate, and that lactate metabolism shapes neuronal activity in the neocortex through K_ATP_ channels. Our results highlight the importance of metabolic crosstalk between neurons and astrocytes for brain function.

**Highlights:**

- Most cortical neurons subtypes express pancreatic beta-cell like K_ATP_ channels.
- Lactate enhances spiking activity via its uptake and closure of K_ATP_ channels.
- Cortical neurons take up and oxidize lactate.
- Cortical neurons produce ATP mainly by oxidative phosphorylation.

## Introduction

The human brain represents 2% of the body mass, yet it consumes about 20% of blood oxygen and glucose which are mandatory energy substrates (Clarke and Sokoloff, 1999). The majority (∼50-80%) of the cerebral energy metabolism is believed to be consumed by the Na^+^/K^+^ ATPase pump to maintain cellular ionic gradients dissipated during synaptic transmission and action potentials (Attwell and Laughlin, 2001;Lennie, 2003). Synaptic and spiking activities are also coupled with local cerebral blood flow and glucose uptake (Devor et al., 2008;Logothetis, 2008). This process, referred to as neurovascular and neurometabolic coupling, is the physiological basis of brain imaging techniques (Raichle and Mintun, 2006) and maintains extracellular glucose within a physiological range of 2-3 mM (Silver and Erecinska, 1994;Hu and Wilson, 1997b). Also, following increased neuronal activity extracellular lactate increases (Prichard et al., 1991;Hu and Wilson, 1997a) for several minutes up to twice of its 2-5 mM basal concentration despite oxygen availability (Magistretti and Allaman, 2018).

Based on the observations that various by-products released during glutamatergic transmission stimulate astrocyte glucose uptake, aerobic glycolysis and lactate release (Pellerin and Magistretti, 1994;Voutsinos-Porche et al., 2003;Ruminot et al., 2011;Choi et al., 2012;Sotelo-Hitschfeld et al., 2015;Lerchundi et al., 2015), lactate has been proposed to be shuttled from astrocytes to neurons to meet neuronal energy needs. This hypothesis is supported by the existence of a lactate gradient between astrocytes and neurons (Machler et al., 2016), the preferential use of lactate as an energy substrate in cultured neurons (Bouzier-Sore et al., 2003;Bouzier-Sore et al., 2006), and its ability to support neuronal activity during glucose shortage (Schurr et al., 1988;Rouach et al., 2008;Wyss et al., 2011;Choi et al., 2012). However, the use of different fluorescent glucose analogues to determine whether astrocytes or neurons take up more glucose during sensory-evoked neuronal activity has led to contradicting results (Chuquet et al., 2010;Lundgaard et al., 2015). Furthermore brain slices and *in vivo* evidence have indicated that synaptic and sensory stimulation enhanced neuronal glycolysis and potentially lactate release by neurons (Ivanov et al., 2014;Diaz-Garcia et al., 2017), thereby challenging the astrocyte-neuron lactate shuttle hypothesis. Hence, the relative contribution of glucose and lactate to neuronal ATP synthesis remains unresolved.

ATP-sensitive potassium channels (K_ATP_) act as metabolic sensors controlling various cellular functions (Babenko et al., 1998). Their open probability (P_o_) is regulated by the energy charge of the cell (*i.e.* the ATP/ADP ratio). While ATP mediates a tonic background inhibition of K_ATP_ channels, cytosolic increases of ADP concentrations that occur as a sequel to enhanced energy demands, increase the P_o_ of K_ATP_ channels. In neurons, electrical activity is accompanied by enhanced sodium influx, which in turn activates the Na^+^/K^+^ ATPase. Activity of this pump alters the submembrane ATP/ADP ratio sufficiently to activate K_ATP_ channels (Tanner et al., 2011). The use of fluorescent ATP/ADP biosensors has demonstrated that K_ATP_ channels are activated (P_o_>0.1) when ATP/ADP ratio is ≤ 5 (Tantama et al., 2013).

K_ATP_ channels are heterooctamers composed of four inwardly rectifying K^+^ channel subunits, Kir6.1 or Kir6.2, and four sulfonylurea receptors, SUR1 or SUR2, the later existing in two splice variants (SUR2A and SUR2B) (Sakura et al., 1995;Aguilar-Bryan et al., 1995;Inagaki et al., 1995b;Isomoto et al., 1996;Inagaki et al., 1996;Chutkow et al., 1996;Yamada et al., 1997;Li et al., 2017;Martin et al., 2017;Lee et al., 2017;Puljung, 2018). The composition in K_ATP_ channel subunits confers different functional properties, pharmacological profiles as well as metabolic sensitivities (Isomoto et al., 1996;Inagaki et al., 1996;Gribble et al., 1997;Yamada et al., 1997;Okuyama et al., 1998;Liss et al., 1999). K_ATP_ channel subunits are expressed in the neocortex (Ashford et al., 1988;Karschin et al., 1997;Dunn-Meynell et al., 1998;Thomzig et al., 2005;Cahoy et al., 2008;Zeisel et al., 2015;Tasic et al., 2016) and have been shown to protect cortical neurons from ischemic injury (Heron-Milhavet et al., 2004;Sun et al., 2006) and to modulate their excitability (Gimenez-Cassina et al., 2012) and intrinsic firing activity (Lemak et al., 2014). K_ATP_ channels could thus be leveraged to decipher electrophysiologically the relative contribution of glucose and lactate to neuronal ATP synthesis. Here, we apply single-cell RT-PCR (scRT-PCR) to identify the mRNA subunit composition of K_ATP_ channel across different neocortical neuron subtypes and demonstrate lactate as the preferred energy substrate that also enhances firing activity.

## Results

### Expression of K_ATP_ channel subunits in identified cortical neurons

We first sought to determine whether K_ATP_ channel subunits were expressed in different neuronal subtypes from the neocortex. Neurons (n=277) of the juvenile rat barrel cortex from layers I to IV (Table S1) were functionally and molecularly characterized in acute slices by scRT-PCR (Figure 1), whose sensitivity was validated from 500 pg of total cortical RNAs (Figure S1A). Neurons were segregated into 7 different subtypes according to their overall molecular and electrophysiological similarity (Figure 1A) using unsupervised Ward’s clustering (Ward, 1963), an approach we previously successfully used to classify cortical neurons (Cauli et al., 2000;Gallopin et al., 2006;Karagiannis et al., 2009). Regular spiking (RS, n=63) and intrinsically bursting (IB, n=10) cells exhibited the molecular characteristics of glutamatergic neurons, with very high single-cell detection rate (n=69 of 73, 95%) of vesicular glutamate transporter 1 (vGluT1) and low detection rate (n=7 of 73, 10%) of glutamic acid decarboxylases (GADs, Figure 1B-E and Table S2), the GABA synthesizing enzymes. This group of glutamatergic neurons distinctly displayed hyperpolarized resting membrane potential (−81.2 ± 0.8 mV), possessed a large membrane capacitance (108.6 ± 3.6 pF), discharged with wide action potentials (1.4 ± 0.0 ms) followed by medium afterhyperpolarizations (mAHs). These neurons did sustain only low maximal frequencies (35.4 ± 1.6 Hz) and showed complex spike amplitude accommodation (Table S3-7). In contrast to RS neurons, IB neurons were more prominent in deeper layers (Table S1) and their bursting activity affected their adaptation amplitudes and kinetics (Figure 1C and Tables S4-5), spike broadening (Figure 1C and Tables S6) and the shape of mAHs (Figure 1C and Tables S7).

**Figure 1.**
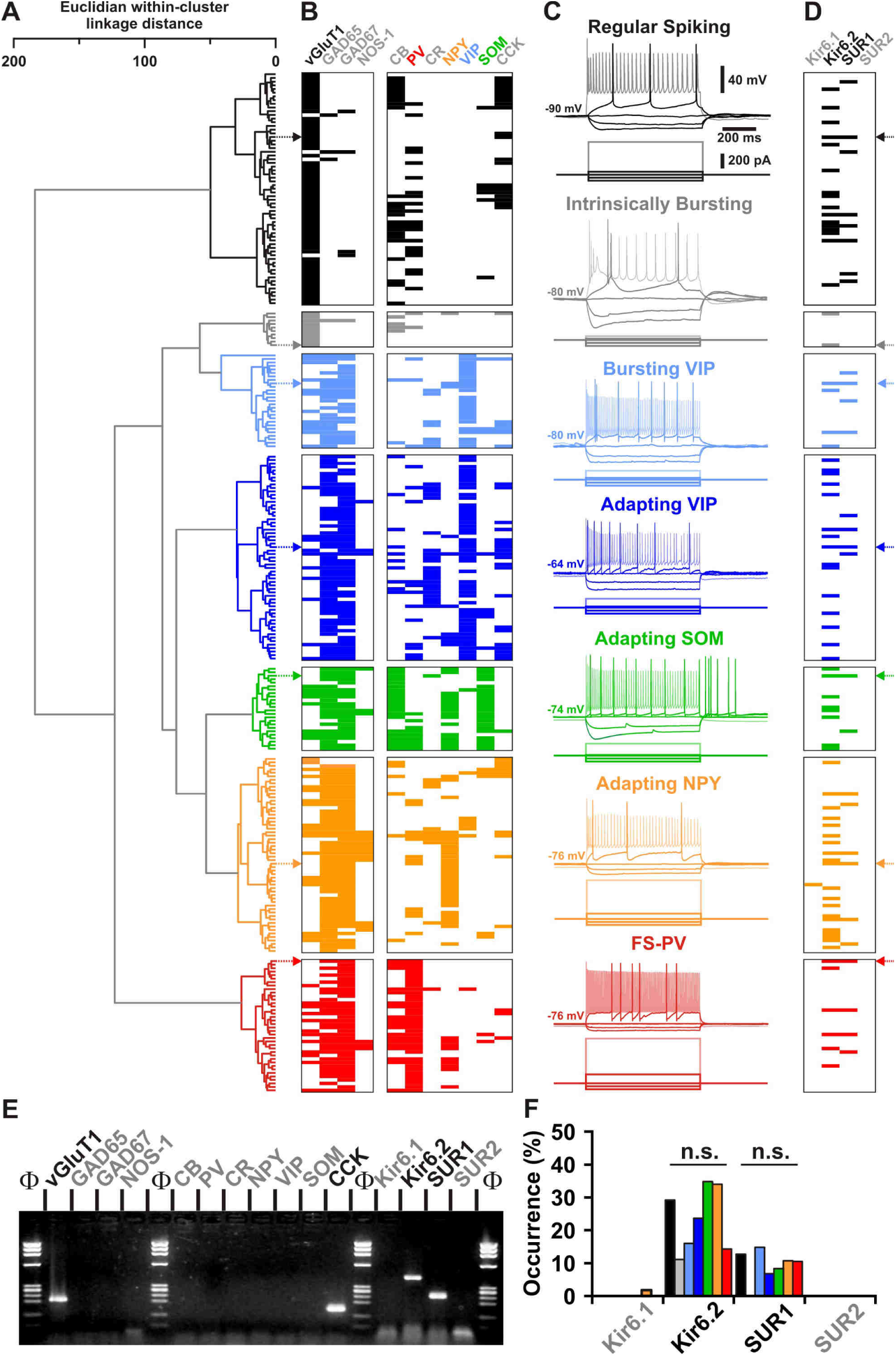
Cortical neuron subtypes express Kir6.2 and SUR1 K_ATP_ channel subunits. (A) Ward’s clustering of 277 cortical neurons (left panel). The x axis represents the average within-cluster linkage distance, and the y axis the individuals. (B) Heatmap of gene expression across the different cell clusters. For each cell, colored and white rectangles indicate presence and absence of genes, respectively. (C) Representative voltage responses induced by injection of current pulses (bottom traces) corresponding to −100, −50 and 0 pA, rheobase and intensity inducing a saturating firing frequency (shaded traces) of a Regular Spiking neuron (black), an Intrinsically Bursting neuron (gray), a Bursting VIP interneuron (light blue), an Adapting VIP interneuron (blue), an Adapting SOM interneuron (green), an Adapting NPY interneuron (orange), and a Fast Spiking-Parvalbumin interneuron (FS-PV, red). The colored arrows indicate the expression profiles of neurons whose firing pattern is illustrated in (C). (D) Heatmap of the expression of the subunits of the K_ATP_ channels in the different clusters. (E) scRT-PCR analysis of the RS neuron depicted in (A-D). (F) Histograms summarizing the expression profile of K_ATP_ channel subunits in identified neuronal types. n.s. not statistically significant.

All other neuronal subtypes were characterized by a high single-cell detection rate of GAD65 and/or GAD67 mRNA (n=202 of 204, 99%, Figure 1B and Table S2) and therefore likely corresponded to GABAergic interneurons. Among GAD-positive population, neurons frequently expressed mRNA for vasoactive intestinal polypeptide (VIP), and in accordance to their electrophysiological phenotypes, were segregated into Bursting VIP (n=27) and Adapting VIP (n=59) neurons. These VIP interneurons were further characterized by high membrane resistance (581 ± 27 MΩ) and small membrane capacitance (52.7 ± 2.3 pF, Figure 1B-C and Tables S2-3).

Other GABAergic interneurons co-expressed somatostatin (SOM) and calbindin (CB) as well as neuropeptide Y (NPY) to a lesser extent and functionally corresponded to Adapting SOM neurons (n=24, Figure 1B and Table S2). They displayed depolarized resting membrane potential, pronounced voltage sags, low rheobases and pronounced afterdepolarizations (Figure 1C and Table S3-6). In another group of GABAergic adapting interneurons located in superficial layers, mRNA for NPY was detected at a high rate (n=31 of 56, 55%). In these Adapting NPY interneurons mRNA for nitric oxide synthase-1 (NOS-1) was detected at a lower rate (Figure 1B and Tables S1-2). In response to suprathreshold depolarizing current steps, these interneurons showed very little spike frequency adaptation (Figure 1C and Table S4). Finally, parvalbumin (PV) was observed in virtually all neurons of a subpopulation termed Fast Spiking-PV interneurons (FS-PV, n=37 of 38, 97%, Figure 1B and Table S2). In comparison to all other cortical neurons described above, they were characterized by low membrane resistance (201 ± 13 MΩ), fast time constant, high rheobase, very short spikes (0.6 ± 0.0 ms) with sharp fast afterhyperpolarizations (fAHs) and the ability to sustain high firing rates (139.9 ± 6.8 Hz) with little to no frequency adaptation (Figure 1C and Tables S3-7). These data thus identified different neuronal subtypes based on their distinctive electrophysiological and molecular features (Ascoli et al., 2008) confirming our previous classification schemes (Cauli et al., 2000;Gallopin et al., 2006;Karagiannis et al., 2009).

The functional and molecular classification of cortical neurons allowed us to probe for the single-cell expression of mRNA for K_ATP_ channel subunits in well defined subpopulations. We observed consistent expression of Kir6.2 and SUR1 between the different cortical neuronal subtypes (Figure 1D-F). Although the scRT-PCR procedure was shown to be reliable for co-detecting all K_ATP_ channel subunits from low amounts of total cortical RNAs (Figure S1A), the overall single-cell detection rates were low for Kir6.2 (n=63 of 248, 25%) and SUR1 (n=28 of 277, 10%). In accordance to previous single-cell studies (Liss et al., 1999;Zeisel et al., 2015;Tasic et al., 2016), our observation suggests an expression at a low copy number at the single-cell level. Apart from a single Adapting NPY neuron (Figure 1D), where Kir6.1 mRNA was detected, our data set provide evidence that for all neocortical neuron types the pattern of detected K_ATP_ channels mRNA expression suggested the co-expression Kir6.2 and SUR1 subunits. This implies that most cortical neurons might be endowed with a pancreatic beta-cell like K_ATP_ channel, which might operate as a metabolic sensor. We next tested this prediction with functional recordings of K_ATP_ channel-mediated whole-cell currents in neocortical neurons.

### Characterization of K_ATP_ channels in cortical neurons

To assess whether K_ATP_ channels are functional in identified cortical neurons (n=18, Figure 2A), we measured the effects of different K_ATP_ channel modulators on whole-cell currents (Q_(3,18)_=32.665, p=3.8 x 10^−7^, Friedman test) and membrane resistances (Q_(3,18)_=40.933, p=6.8 x 10^−9^). Pinacidil (100 µM), a SUR2-preferring K_ATP_ channel opener (Inagaki et al., 1996;Moreau et al., 2005), had little or no effect on current (4.1 ± 3.7 pA, p=0.478) and membrane resistance (−9.6 ± 3.7 %, p=0.121, Figure 2B-C). By contrast, diazoxide (300 µM), an opener acting on SUR1 and SUR2B-containing K_ATP_ channels (Inagaki et al., 1996;Moreau et al., 2005), consistently induced an outward current (45.0 ± 9.6 pA, p=4.8 x 10^−5^) and a decrease in membrane resistance (−34.5 ± 4.3%, p=3.6 x 10^−5^) indicative of the activation of a hyperpolarizing conductance (Figure 2B-C). The sulfonylurea tolbutamide (500 µM, Figure 2B-C), a K_ATP_ channel blocker (Ammala et al., 1996;Isomoto et al., 1996;Gribble et al., 1997;Isomoto and Kurachi, 1997), did not change whole-cell basal current (−6.6 ± 3.0 pA, p=0.156) or membrane resistance (20.5 ± 7.5%, p=3.89 x 10^−2^). Conversely, tolbutamide dramatically reversed diazoxide effects on both current (p=4.1 x 10^−8^) and membrane resistance (p=5.8 x 10^−10^).

**Figure 2.**
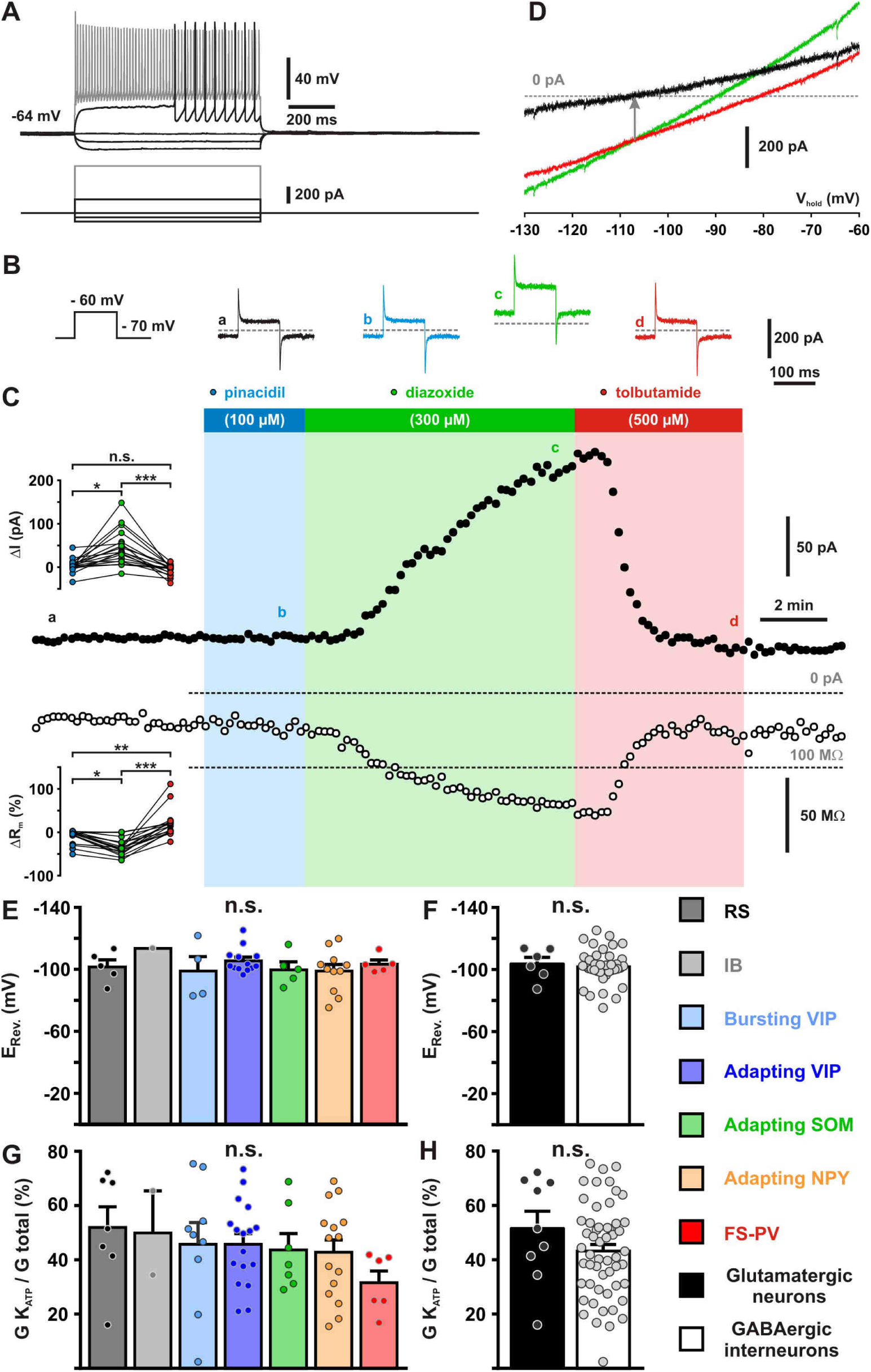
Pharmacological and biophysical characterization of K_ATP_ channels in cortical neurons. (A) Representative voltage responses of a FS-PV interneuron induced by injection of current pulses (bottom traces). (B) Protocol of voltage pulses from −70 to −60 mV (left trace). Responses of whole-cell currents in the FS-PV interneurons shown in (A) in control condition (black) and in presence of pinacidil (blue), piazoxide (green) and tolbutamide (red) at the time indicated by a-d in (C). (C) Stationary currents recorded at −60 mV (filled circles) and membrane resistance (open circles) changes induced by K_ATP_ channel modulators. The colored bars and shaded zones indicate the duration of application of K_ATP_ channel modulators. Upper and lower insets: changes in whole-cell currents and relative changes in membrane resistance induced by K_ATP_ channel modulators, respectively. (D) Whole cell current-voltage relationships measured under diazoxide (green trace) and tolbutamide (red trace). K_ATP_ I/V curve (black trace) obtained by subtracting the curve under diazoxide by the curve under tolbutamide. The arrow indicates the reversal potential of K_ATP_ currents. (E-H) Histograms summarizing the K_ATP_ current reversal potential (E,F) and relative K_ATP_ conductance (G,H) in identified neuronal subtypes (E,G) or between glutamatergic and GABAergic neurons (F,G). Data are expressed as mean ± s.e.m., and the individual data points are depicted. n.s. not statistically significant.

In virtually all neuronal subtypes (H_(6,43)_=2.274, p=0.810, Kruskal–Wallis H test) or groups (t_(42)_=0.3395, p=0.736, Student’s t-test), the diazoxide-tolbutamide current/voltage relationship reversed very close to the theoretical potassium equilibrium potential (E_K_=-106.0 mV, Figure 2D-F) confirming the opening of a selective potassium conductance. Besides its effects on plasma membrane K_ATP_ channels, diazoxide is also a mitochondrial uncoupler (Drose et al., 2006) which increases reactive oxygen species (ROS) production. This might stimulate Ca^2+^ sparks and large-conductance Ca^2+^-activated potassium channels (Xi et al., 2005) leading to potential confounding effects. This possibility was ruled out by the observation that Mn(III)tetrakis(1-methyl-4-pyridyl)porphyrin (MnTMPyP, 25 µM), a ROS scavenger (D’Agostino et al., 2007), did not reduce the diazoxide-tolbutamide responses on current (t_(10)_=0.76559, p=0.462, Figure S2A,B) and conductance (t_(10)_=1.24758, p=0.241, Figure S2A,C).

Cortical neurons exhibited K_ATP_ conductances of similar value between their subtypes (H_(6,63)_=5.6141, p=0.468) or groups (U_(9,54)_=233, p=0.855, Mann–Whitney U test, Figures S3A,B). K_ATP_ channels activated by diazoxide essentially doubled the whole cell conductance in the subthreshold membrane potential compared to control or tolbutamide conditions, regardless of neuronal subtypes (H_(6,63)_=5.4763, p=0.484) or groups (t_(61)_=1.324, p=0.191, Figures 2G,H). Also, K_ATP_ current density was similar (H_(6,63)_=4.4769, p=0.612, U_(9,54)_=240.5, p=0.965, Figures S3C,D). Together with the pinacidil unresponsiveness, these data indicate that the large majority of cortical neurons expressed functional SUR1-mediated K_ATP_ channels in a homogeneous fashion across different subpopulations. To confirm that Kir6.2 is the pore-forming subunit of K_ATP_ channels in cortical neurons, we used a genetic approach based on Kir6.2 knock-out mice (Miki et al., 1998). We first verified the expression of Kir6.2 and SUR1 subunits in pyramidal cells from wild type mice by scRT-PCR (Figure 3A,B). We next used a dialysis approach by recording neurons with an ATP-free pipette solution (Miki et al., 2001) enriched in sodium (20 mM) to stimulate submembrane ATP depletion and ADP production by the Na^+^/K^+^ ATPase, which is known to activate K_ATP_ channels (Figure 3H). We confirmed that ATP1a1 and ATP1a3 (Figure 3B) were the main α-subunits of the Na^+^/K^+^ ATPase pump in pyramidal neurons (Zeisel et al., 2015;Tasic et al., 2016). Dialysis of ATP-free/20 mM Na^+^-pipette solution indeed induced a standing outward current in Kir6.2^+/+^ neurons within 5 minutes (46.7 ± 19.0 pA at −50 mV, n=26) that was reversing close to E_K_ (Figure 3C,E). In contrast, this current was not observed in Kir6.2^-/-^ neurons (−59.9 ± 11.9 pA, n=22, U_(26,22)_=78, p=2.4221 x 10^−6^, one-tailed, Figure 3D-G). Collectively, these data show that cortical neurons predominately express functional K_ATP_ channels composed of Kir6.2 and SUR1 subunits.

**Figure 3.**
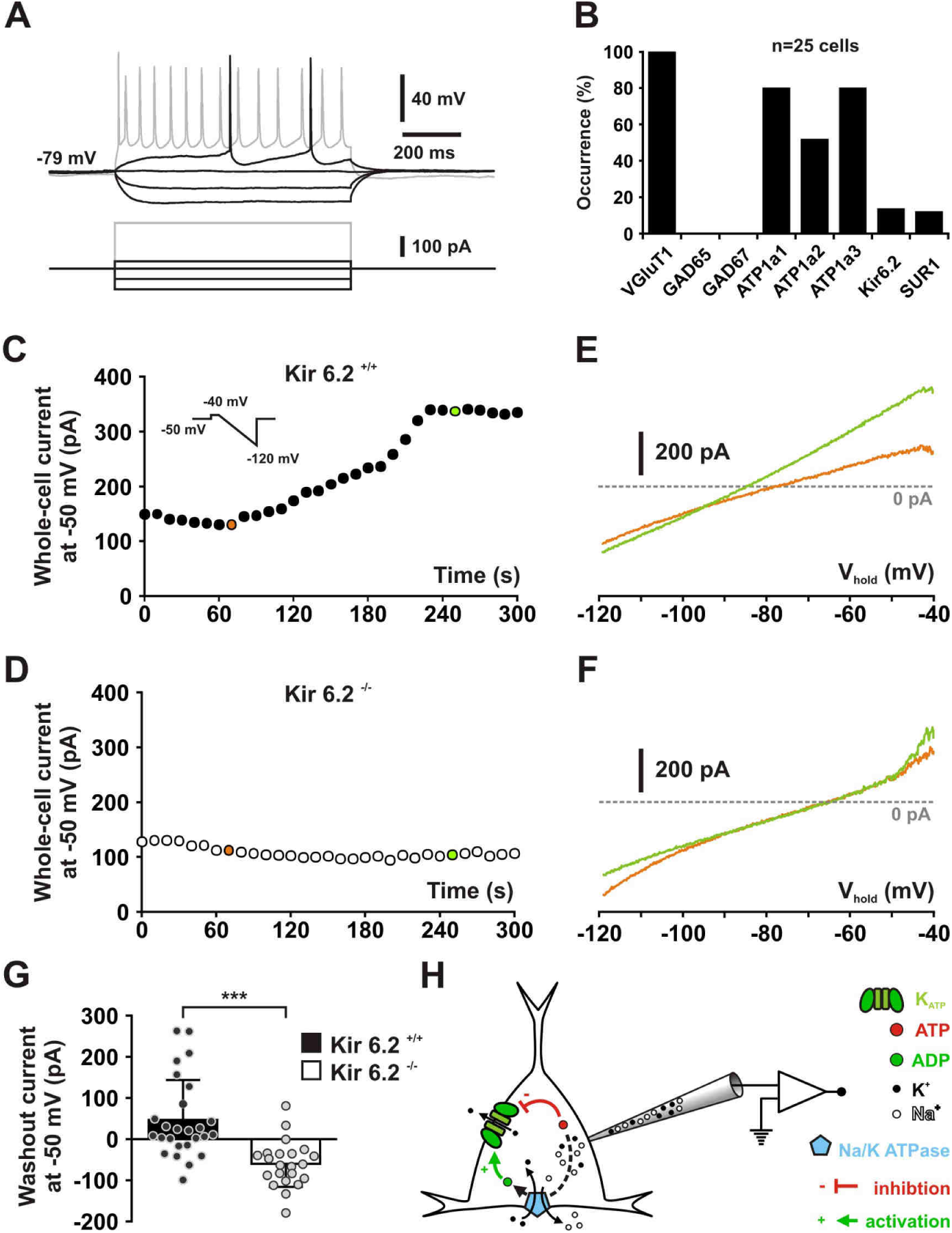
Kir6.2 is the pore forming subunit of K_ATP_ channels in cortical neurons. (A) Representative voltage responses of a mouse layer II/III RS pyramidal cell induced by injection of current pulses (bottom traces). (B) Histograms summarizing the expression profile of vGluT1, GAD65 and 67, the ATP1a1-3 subunits of the Na/K ATPase and the Kir6.2 and SUR1 K_ATP_ channel subunits in layer II/III RS pyramidal cells from Kir6.2^+/+^ mice. (C, D) Whole-cell stationary currents recorded at −50 mV during dialysis with ATP-free pipette solution in RS neurons of Kir6.2^+/+^ (C) and Kir6.2^-/-^ (D) mice. Inset; voltage clamp protocol. (E, F) Current-voltage relationships obtained during ATP washout at the time indicated by green and orange circles in (C, D) in RS neurons of Kir6.2^+/+^ (E) and Kir6.2^-/-^ (F) mice. (G) Histograms summarizing the whole-cell ATP washout currents in Kir6.2^+/+^ (black) and Kir6.2^-/-^ (white) RS neurons. Data are expressed as mean ± s.e.m., and the individual data points are depicted. (H) Diagram depicting the principle of the ATP washout experiment.

### Modulation of neuronal excitability and activity by K_ATP_ channel

Despite their large diversity, cortical neurons display a homogenous functional expression of K_ATP_ channels, questioning how these channels integrate the metabolic environment to adjust neuronal activity. To address this question, we first evaluated in identified cortical neurons (n=39) the ability of K_ATP_ channels to modulate neuronal excitability, notably by measuring membrane potentials (Q_(2,39)_=38.000, p=5.6 x 10^−9^) and membrane resistances (Q_(2,39)_=40.205, p=1.9 x 10^−9^), as well as spiking activity (Q_(2,39)_=28.593, p=6.2 x 10^−9^). Following electrophysiological identification, the K_ATP_ channel blocker tolbutamide was applied, which resulted in a slight depolarization (ΔV_m_=2.6 ± 0.8 mV, p=1.74 x 10^−2^, Figure 4A,D) and increase in membrane resistance (ΔR_m_=78 ± 31 MΩ, p=1.52 x 10^−3^, Figure 4B,E). These effects were strong enough to trigger and stimulate the firing of action potentials (ΔF=0.3 ± 0.1 Hz, p= 9.21 x 10^−3^, Figure 4A,C,F). By contrast, diazoxide hyperpolarized cortical neurons (−4.0 ± 0.6 mV, p=1.87 x 10^−4^, Figure 4A,D), decreased their membrane resistance (−39 ± 23 MΩ, p=1.52 x 10^−3^, Figure 4B,E) but did alter their rather silent basal spiking activity (−0.1 ± 0.1 Hz, p=0.821, Figure 4A,C,F).

**Figure 4.**
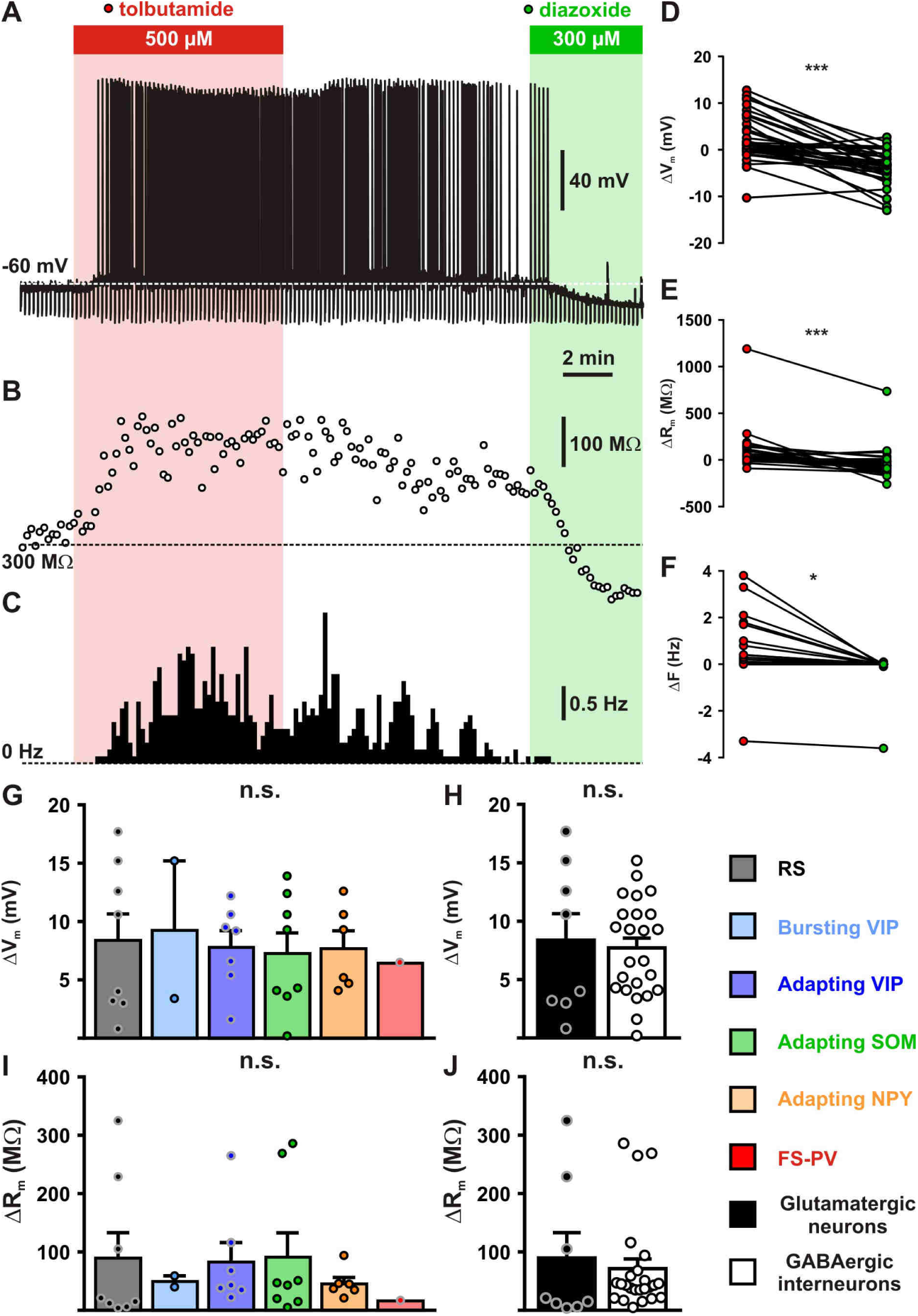
Modulation of cortical neuronal excitability and activity by K_ATP_ channels. (A-C) Representative example showing the changes in membrane potential (A), resistance (B, open circles) and spiking activity (C) induced by application of tolbutamide (red) and diazoxide (green). The colored bars and shaded zones indicate the application duration of K_ATP_ channel modulators. (D-F) Relative changes in membrane potential (D), resistance (E) and firing rate (F) induced by tolbutamide and diazoxide in cortical neurons. (G-J) Histograms summarizing the modulation of membrane potential (G, H_(5,32)_=0.14854, p=0.999, and H, U_(8,24)_=95.5, p=0.991) and resistance (I, H_(5,32)_=3.00656, p=0.699, and J, U_(8,24)_=73, p=0.329) by K_ATP_ channels in neuronal subtypes (G, I) and groups (H, J). Data are expressed as mean ± s.e.m., and the individual data points are depicted. n.s. not statistically significant.

Most cortical neurons (n=32 of 39) showed modulation of neuronal excitability by both K_ATP_ channel modulators and were considered to be responsive. A similar proportion of responsive neurons was observed between neuronal subtypes (Figure S4A, X^2^_(5)_=7.313, p=0.1984) or groups (Figure S4B, p=0.9999, Fisher’s exact test). The apparent relative lack of responsiveness in FS-PV interneurons (Figure S4A), despite a whole-cell K_ATP_ conductance similar to that of other neuronal types (Figure S3A), is likely attributable to their low input resistance (Table S3) making K_ATP_ channels less effective to change membrane potential. Overall, K_ATP_ channels modulated membrane potential, resistance and firing rate by up to 7.6 ± 0.9 mV, 76 ± 17% and 0.5 ± 0.2 Hz, respectively. This modulation of neuronal excitability (Figure 4G-J) and activity (Figure S4C,D) was similar between neuronal subtypes or groups (Figure 4H-J and S4C-E). Thus, K_ATP_ channels modulate the excitability and activity of all subtypes of cortical neurons.

### Enhancement of neuronal activity by lactate via modulation of K_ATP_ channels

The uniform expression of metabolically highly sensitive K_ATP_ channels by cortical neurons suggests their ability to couple the local glycolysis capacity of astrocytes with spiking activity. We therefore evaluated whether extracellular changes in glucose and lactate could differentially shape the spiking activity of cortical neurons through their energy metabolism and K_ATP_ channel modulation. Importantly, to preserve intracellular metabolism, neurons were recorded in perforated patch-configuration. Stable firing rates of about 4 Hz inducing ATP consumption by the Na^+^/K^+^ ATPase (Attwell and Laughlin, 2001) were evoked by applying a depolarizing current and continuously monitored throughout changes in extracellular medium (Figure 5A, Q_(2,16)_=22.625, p=1.222 x 10^−5^).

**Figure 5.**
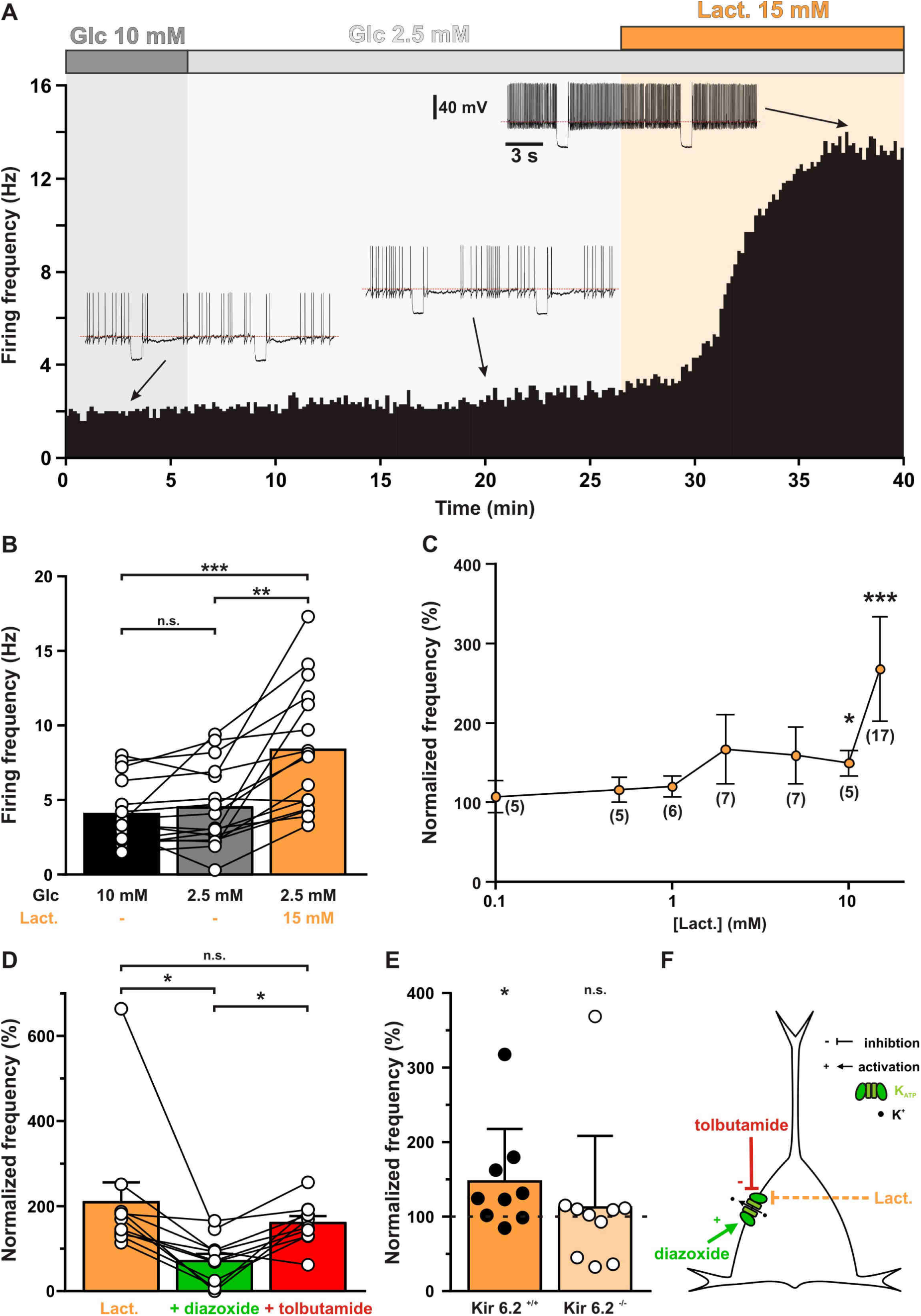
Lactate enhances cortical neuronal activity via K_ATP_ channel modulation. (A) Representative perforated patch recording of an adapting VIP neuron showing the modulation of firing frequency induced by changes in the extracellular concentrations of metabolites. The colored bars and shaded zones indicate the concentration in glucose (grey) and lactate (orange). Voltage responses recorded at the time indicated by arrows. The red dashed lines indicate −40 mV. (B) Histograms summarizing the mean firing frequency during changes in extracellular concentration of glucose (black and grey) and lactate (orange). Data are expressed as mean ± s.e.m., and the individual data points are depicted. n.s. not statistically significant. (C) Dose-dependent enhancement of firing frequency by lactate. Data are normalized by the mean firing frequency in absence of lactate and are expressed as mean ± s.e.m. Numbers in brackets indicate the number of recorded neurons at different lactate concentrations. (D) Histograms summarizing the normalized frequency under 15 mM lactate (orange) and its modulation by addition of diazoxide (green) or tolbutamide (red). Data are expressed as mean ± s.e.m., and the individual data points are depicted. n.s. not statistically significant. (E) Histograms summarizing the enhancement of normalized frequency by 15 mM lactate in Kir6.2 ^+/+^ (orange) and Kir6.2 ^-/-^ (pale orange) mouse cortical neurons. The dash line indicates the normalized mean firing frequency in absence of lactate. Data are expressed as mean ± s.e.m., and the individual data points are depicted. (F) Diagram depicting the enhancement of neuronal activity by lactate via modulation of K_ATP_ channels.

Decreasing extracellular glucose from 10 mM to a normoglycemic concentration of 2.5 mM (Silver and Erecinska, 1994;Hu and Wilson, 1997b) did not change firing rate (Figure 5A,B, p=0.2159) of cortical neurons (n=16). By contrast, supplementing extracellular 2.5 mM glucose with 15 mM lactate, a concentration observed in the blood after an intense physical exercise (Quistorff et al., 2008) and corresponding to the same number of carbon atoms as 10 mM glucose, roughly doubled the firing rate compared to both 2.5 (p=7.829 x 10^−4^) and 10 mM glucose (p=4.303 x 10^−6^) conditions. Firing rate enhancement by lactate was dose-dependent (H_(8,76)_= 35.142, p= 1.052 x 10^−5^) and reached statistical significance above 5 mM (Figure 5C). We reasoned that this effect could be mediated by K_ATP_ channel closure. Indeed, the increase in firing rate by lactate (209 ± 49 %) was strongly reduced by the K_ATP_ channel activator diazoxide (71 ± 18 %, p=3.346 x 10^−3^, Figure 5D). Tolbutamide reversed diazoxide’s effect (160 ± 17 %, p=9.345 x 10^−3^) but did not increase firing rate further (p=0.5076). This occlusion of tolbutamide’s effect by 15 mM lactate also suggests that this concentration reaches saturating levels and is the highest metabolic state that can be sensed by K_ATP_ channels. Enhancement of neuronal activity by lactate was also observed in Kir6.2^+/+^ cortical neurons (147 ± 25 %, p=2.840 x 10^−2^) but not in Kir6.2^-/-^ mice (112 ± 32 %, p=0.8785, Figure 5E). These observations indicate that lactate enhances neuronal activity via a closure of K_ATP_ channels (Figure 5F).

### Mechanism of lactate-sensing

To determine whether lactate-sensing involves intracellular lactate oxidative metabolism and/or extracellular activation of the lactate receptor GPR81, we next probed the expression of monocarboxylate transporters (MCT), which allow lactate uptake. Consistent with mouse RNAseq data (Zeisel et al., 2015;Tasic et al., 2016), MCT1 and MCT2 were the main transporters detected in rat cortical neurons by scRT-PCR, although with relatively low single cell detection rates (54 of 277, 19.5% and 78 of 277, 28.2%, for MCT1 and 2, respectively, Figure 6A and S5).

**Figure 6.**
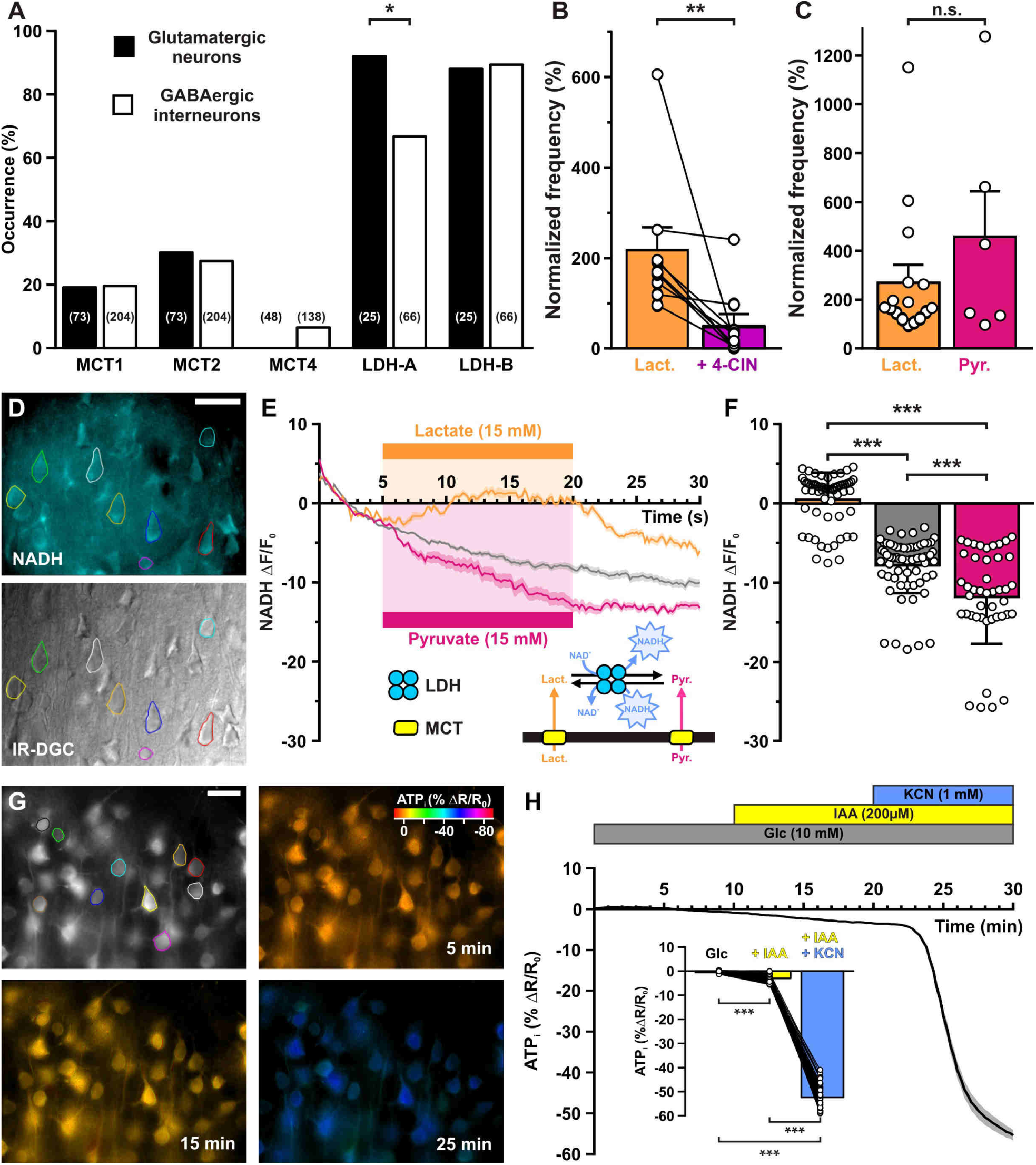
Lactate enhancement of cortical neuronal activity involves lactate uptake and metabolism. (A) Histograms summarizing the expression profile of the monocarboxylate transporters MCT1, 2 and 4 and LDH A and B lactate dehydrogenase subunits in glutamatergic neurons (black) and GABAergic interneurons (white). The numbers in brackets indicate the number of analyzed cells. (B) Histograms summarizing the enhancement of normalized frequency by 15 mM lactate (orange) and its suppression by the MCTs inhibitor 4-CIN (purple). Data are expressed as mean ± s.e.m., and the individual data points are depicted. (C) Histograms summarizing the enhancement of normalized frequency by 15 mM lactate (orange) and pyruvate (magenta). Data are expressed as mean ± s.e.m., and the individual data points are depicted n.s. not statistically significant. (D) Widefield NADH autofluorescence (upper panel, scale bar: 20 µm) and corresponding field of view observed under IR-DGC (lower panel). The somatic regions of interest are delineated. (E) Mean relative changes in NADH autofluorescence in control condition (grey) and in response to 15 mM lactate (orange) or pyruvate (magenta). The colored bars indicate the duration of applications. Data are expressed as mean ± s.e.m. Inset: diagram depicting the NADH changes induced by lactate and pyruvate uptake by MCT and their interconversion by LDH. (F) Histograms summarizing the mean relative changes in NAD(P)H autofluorescence measured during the last 5 minutes of 15 mM lactate (orange) or pyruvate (magenta) application and corresponding time in control condition (grey). Data are expressed as mean ± s.e.m., and the individual data points are depicted. (G) Widefield YFP fluorescence of the ATP biosensor AT1.03^YEMK^ (upper left panel, scale bar: 30 µm) and pseudocolor images showing the intracellular ATP (YFP/CFP ratio value coded by pixel hue, see scale bar in upper right panel) and the fluorescence intensity (coded by pixel intensity) at different times under 10 mM extracellular glucose (upper right panel) and after addition of IAA (lower left panel) and KCN (lower right panel). (H) Mean relative changes in intracellular ATP (relative YFP/CFP ratio) measured under 10 mM extracellular glucose (grey) and after addition of IAA (yellow) and KCN (blue). Data are expressed as mean ± s.e.m. The colored bars indicate the time and duration of metabolic inhibitor application. Inset: Histograms summarizing the mean relative changes in intracellular ATP (relative YFP/CFP ratio) ratio under 10 mM extracellular glucose (grey) and after addition of IAA (yellow) and KCN (blue). Data are expressed as mean ± s.e.m., and the individual data points are depicted.

The expression of monocarboxylate transporters in cortical neurons is compatible with lactate uptake and metabolism leading to the closure of K_ATP_ channels and an increase in firing rate. We thus evaluated whether lactate uptake was needed for lactate-sensing. We used 250 µM α-cyano-4-hydroxycinnamic acid (4-CIN), a concentration blocking lactate uptake while only moderately altering mitochondrial pyruvate carrier in brain slices (Schurr et al., 1999;Ogawa et al., 2005;Galeffi et al., 2007). 4-CIN reversed the increased firing rate induced by lactate (Figure 6B, T(9)=0, p=7.686 x 10^−3^) indicating that facilitated lactate transport is required for K_ATP_ channel closure and in turn firing rate acceleration.

A mechanism of lactate-sensing involving an intracellular lactate oxidative metabolism would also require the expression of lactate dehydrogenase (LDH), that reversibly converts lactate and nicotinamide adenine dinucleotide (NAD^+^) to pyruvate and NADH (Figure 6E, inset). We thus also probed for the expression of LDH subunits. LDH-A and LDH-B were observed in a large majority of cortical neurons with LDH-A being more frequent in glutamatergic neurons than in GABAergic interneurons (p=1.61 x 10^−2^, Figure 6A and S5). Nonetheless, neuron subtypes analysis did not allow to disclose which populations express less frequently LDH-A. (Figure S5). To confirm the ability of cortical neurons to take up and oxidize lactate we also visualized NADH fluorescence dynamics (Chance et al., 1962) induced by bath application of lactate. Widefield somatic NADH fluorescence appeared as a diffuse labeling surrounding presumptive nuclei (Figure 6D). Consistent with lactate transport by MCTs and oxidization by LDH, NADH was increased under lactate application (U_(61,67)_=196, p=1.2 10^−18^, Figure 6E-F).

Since the lactate receptor GPR81 has been observed in the cerebral cortex (Lauritzen et al., 2014), lactate-sensing might also involve this receptor. This possibility was ruled out by the observation that pyruvate (15 mM), which is transported by MCTs (Broer et al., 1998;Broer et al., 1999) but does not activate GPR81 (Ahmed et al., 2010), enhanced firing rate to an extent similar to that of lactate (Figure 6C, U_(16,6)_=43, p=0.7468). In line with its uptake and reduction, pyruvate also decreased NADH (Figure 6E-F, U_(44,67)_=868, p=2.08 x 10^−4^).

The requirement of monocarboxylate transport and the similar effect of lactate and pyruvate on neuronal activity suggest that once taken up, lactate would be oxidized into pyruvate and metabolized by mitochondria to produce ATP, leading in turn to a closure of K_ATP_ channels and increased firing rate. The apparent absence of glucose responsiveness in cortical neurons also suggests that glycolysis contributes modestly to ATP production. To determine the relative contribution of glycolysis and oxidative phosphorylation to ATP synthesis, we transduced the genetically encoded fluorescence resonance energy transfer (FRET)-based ATP biosensor AT1.03^YEMK^ (Imamura et al., 2009) using a recombinant Sindbis virus. AT1.03^YEMK^ fluorescence was mostly observed in pyramidal shaped cells (Figure 6G), consistent with the strong tropism of this viral vector towards pyramidal neurons (Piquet et al., 2018). Blocking glycolysis with 200 µM iodoactic acid (IAA) decreased modestly the FRET ratio by 2.9 ± 0.2% (Figure 6H, p=2.55 x 10^−9^). By contrast, adding potassium cyanide (KCN, 1mM), a respiratory chain blocker, reduced the FRET ratio to a much larger extent (52.3 ± 0.6%, Figure 6H, p=2.55 x 10^−9^). KCN also induced a strong NADH fluorescence increase (Figure S6A-B, U_(12,42)_=0, p=5.83 x 10^−12^), indicating a highly active oxidative phosphorylation in cortical neurons.

## Discussion

We report that extracellular lactate and pyruvate, but not glucose, enhance the activity of cortical neurons through a mechanism involving facilitated transport and the subsequent closure of K_ATP_ channels composed of Kir6.2 and SUR1 subunits. ATP synthesis derives mostly from oxidative phosphorylation and weakly from glycolysis in cortical neurons. Together with their ability to oxidize lactate by LDH, these observations suggest that lactate is a preferred energy substrate over glucose in cortical neurons. Besides its metabolic importance lactate also appears as a signaling molecule enhancing firing activity (Figure 7). This suggests that an efficient neurovascular and neurometabolic coupling could define a time window of an up state of lactate during which neuronal activity and plasticity would be locally enhanced (Suzuki et al., 2011;Jimenez-Blasco et al., 2020).

**Figure 7.**
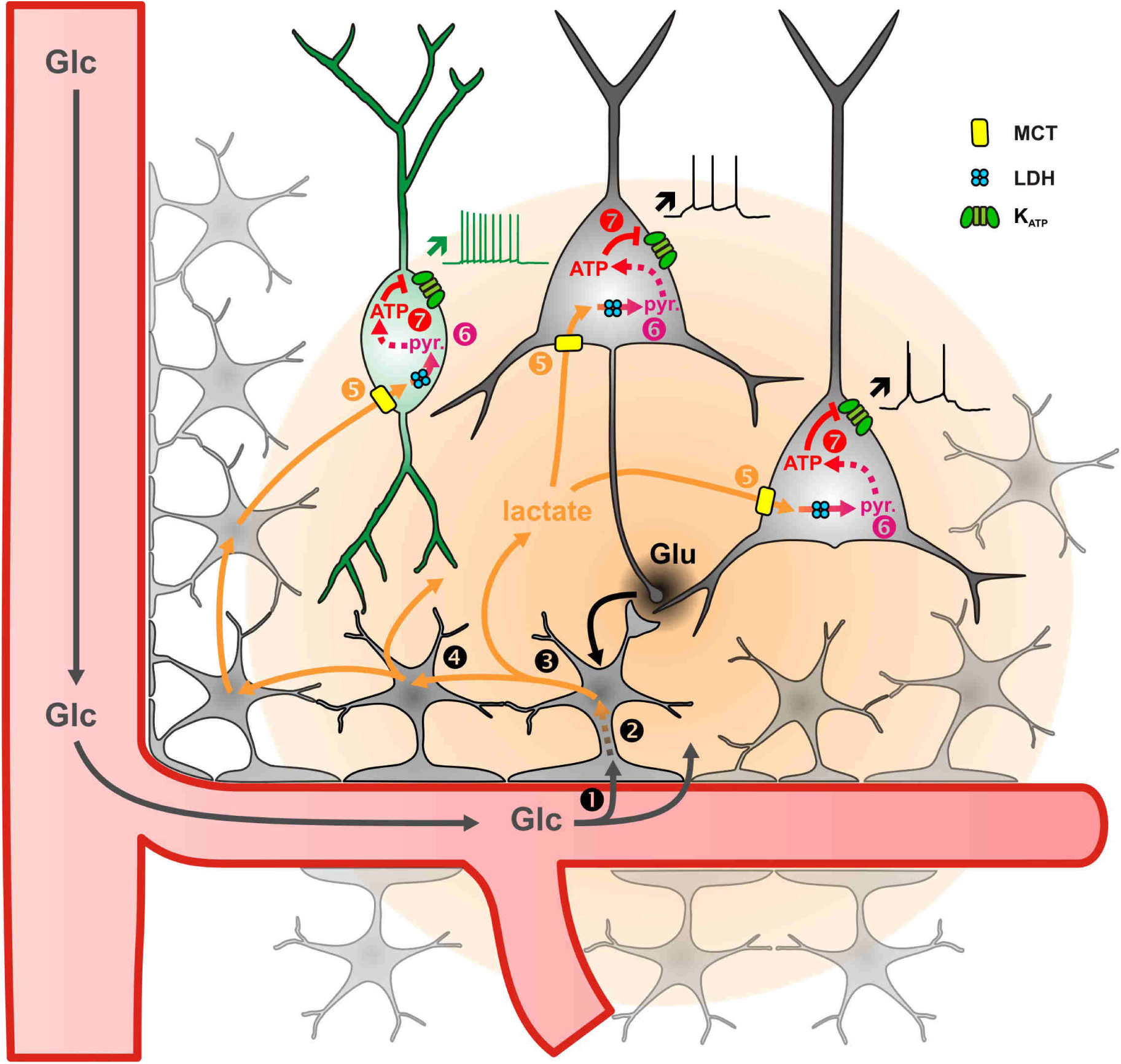
Diagram summarizing the mechanism of lactate-sensing in the cortical network. Glutamate (Glu) released during synaptic transmission stimulates ❶ blood glucose (Glc) uptake in astrocytes, ❷ aerobic glycolysis, ❸ lactate release and ❹ diffusion through the astrocytic network. Lactate is then ❺ taken up by neurons via monobarboxylate transporters (MCT) and ❻ oxidized into pyruvate by lactate dehydrogenase (LDH). The ATP produced by pyruvate oxidative metabolism ❼ closes K_ATP_ channels and increases the spiking activity of both pyramidal cells (black) and inhibitory interneurons (green). The color gradient of the circles represents the extent of glutamate (black) and lactate (orange) diffusion, respectively. Dashed arrows indicate multisteps reactions.

### K_ATP_ channel subunits in cortical neurons

Similarly to neurons of the hippocampal formation (Zawar et al., 1999;Cunningham et al., 2006;Sada et al., 2015) we found that, regardless of the neuronal type, most neocortical neurons express diazoxide-sensitive, but pinacidil-insensitive K_ATP_ channels (Cao et al., 2009). In agreement with this pharmacological profile (Inagaki et al., 1996) and the absence of functional K_ATP_ channels in Kir6.2^-/-^ neurons, we observed Kir6.2 and SUR1 subunits as the main components of K_ATP_ channels. Their low detection rate is presumably due to the low copy number of their mRNAs, as described in single cortical neurons from mice (Zeisel et al., 2015;Tasic et al., 2016), and making their observation by scRT-PCR challenging (Tsuzuki et al., 2001). Since Kir6.2 is an intronless gene, collection of the nucleus was avoided to prevent potential false positives. Thus, neurons positive for both Kir6.2 and somatostatin intron, taken as an indicator of genomic DNA (Hill et al., 2007;Devienne et al., 2018), were discarded from Kir6.2 expression analysis. Unavoidably, this procedure does reduce the amount of cytoplasm collected, thereby decreasing the detection rate of both Kir6.2 and SUR1.

Consistent with the preferred expression of Kir6.1 in mural and endothelial cells (Bondjers et al., 2006;Zeisel et al., 2015;Tasic et al., 2016;Aziz et al., 2017;Vanlandewijck et al., 2018;Saunders et al., 2018), this subunit was only observed in one out of 277 cortical neurons analyzed. Similarly, SUR2B, the SUR2 variant expressed in forebrain (Isomoto et al., 1996) and cortex (Figure S1), whose presence is largely restricted to vascular cells (Zeisel et al., 2015), was not observed in cortical neurons.

### Relative sensitivity of cortical neurons to glucose, lactate and pyruvate

Consistent with previous observations (Yang et al., 1999), decreasing extracellular glucose from standard slice concentrations down to a normoglycemic level did not alter firing rates of cortical neurons. However, their activity is silenced during hypoglycemic episodes through K_ATP_ channels activation (Yang et al., 1999;Zawar and Neumcke, 2000;Molnar et al., 2014;Sada et al., 2015). This relative glucose unresponsiveness is in contrast with pancreatic beta cells and hypothalamic glucose-excited neurons whose activity is regulated over a wider range of glucose concentrations by K_ATP_ channels also composed with Kir6.2 and SUR1 subunits (Aguilar-Bryan et al., 1995;Inagaki et al., 1995a;Miki et al., 1998;Yang et al., 1999;Miki et al., 2001;Tarasov et al., 2006;Varin et al., 2015). The inability of cortical neurons to regulate their spiking activity at glucose levels beyond normoglycemia is likely due to the lack of glucokinase, a hexokinase which catalyzes the first step of glycolysis and acts as a glucose sensor in the millimolar range (German, 1993;Yang et al., 1999). As earlier reported, hexokinase-1 (HK1) is the major isoform in cortical neurons (Zeisel et al., 2015;Tasic et al., 2016;Piquet et al., 2018). Since this enzyme has a micromolar affinity for glucose and is inhibited by its product, glucose-6-phosphate (Wilson, 2003), HK1 is likely already saturated and/or inhibited during normoglycemia thereby limiting glycolysis. Nonetheless, HK1 saturation/inhibition can be mitigated when energy consumption is high (Attwell and Laughlin, 2001;Wilson, 2003;Tantama et al., 2013), and then glucose can probably modulate neuronal activity via a high affinity mechanism, as evidenced by slow oscillations of spiking activity involving synaptic transmission (Cunningham et al., 2006) or by the use of glucose-free whole-cell patch-clamp solution (Kawamura, Jr. et al., 2010) that mimics high glucose consumption (Piquet et al., 2018;Diaz-Garcia et al., 2019).

Similarly to glucose-excited hypothalamic neurons (Yang et al., 1999;Song and Routh, 2005), but in contrast with pancreatic beta cells (Newgard and McGarry, 1995), cortical neurons were dose-dependently excited by lactate. This lactate sensitivity is consistent with lactate transport and oxidization in hypothalamic and cortical neurons (Ainscow et al., 2002;Sada et al., 2015;Diaz-Garcia et al., 2017) which are low in beta cells (Sekine et al., 1994;Pullen et al., 2011). Pyruvate had a similar effect to lactate in cortical neurons under normoglycemic condition whereas it only maintains the activity of hypothalamic glucose-excited neurons during hypoglycemia (Yang et al., 1999) and barely activates pancreatic beta cells (Dufer et al., 2002). Thus, cortical neurons display a peculiar metabolic sensitivity to monocarboxylates. Our data also suggest that under normoglycemic conditions a portion of K_ATP_ channels are open when cortical neurons fire action potentials.

### Mechanism of lactate-sensing

Our pharmacological, molecular and genetic evidence indicates that the closure of K_ATP_ channels is responsible for the firing rate enhancement by lactate. Since K_ATP_ channels can be modulated by G protein-coupled receptors (Kawamura, Jr. et al., 2010), lactate-sensing might have been mediated by GPR81, a G_i_ protein-coupled lactate receptor expressed in the cerebral cortex (Lauritzen et al., 2014). This possibility is however unlikely since the activation of GPR81 inhibits cultured cortical neurons (Bozzo et al., 2013;de Castro Abrantes H. et al., 2019) and we show here that enhancing effect pyruvate on neuronal activity was similar to that of lactate, although pyruvate does not activate GPR81 (Ahmed et al., 2010).

We found that lactate-sensing was critically dependent on lactate transport and we confirmed the capacity of cortical neurons to take up and oxidize lactate (Bittar et al., 1996;Laughton et al., 2000;Bouzier-Sore et al., 2003;Wyss et al., 2011;Choi et al., 2012;Sada et al., 2015;Machler et al., 2016). LDH metabolites, including pyruvate and oxaloacetate, can lead to K_ATP_ channel closure (Dhar-Chowdhury et al., 2005;Sada et al., 2015) and could mediate lactate-sensing. An intermediate role of oxaloacetate in lactate-sensing is compatible with enhanced Krebs cycle and oxidative phosphorylation, which leads to an increased ATP/ADP ratio and the closure of K_ATP_ channels (Figure 7). In contrast to oxaloacetate, intracellular ATP was found to be ineffective for reverting K_ATP_ channel opening induced by LDH inhibition (Sada et al., 2015). Interestingly, hippocampal interneurons were found to be insensitive to glucose deprivation in whole cell configuration (Sada et al., 2015) but not in perforated patch configuration (Zawar and Neumcke, 2000) whereas almost the opposite was found in CA1 pyramidal cells. Whether altered intracellular metabolism by whole-cell recording accounted for the apparent lack of ATP sensitivity remains to be determined.

### Lactate as an energy substrate for neurons and an enhancer of spiking activity and neuronal plasticity

We confirmed that the ATP produced by cortical neurons was mostly derived from oxidative phosphorylation and marginally from glycolysis (Almeida et al., 2001;Hall et al., 2012). Together with the enhancement of spiking activity through K_ATP_ channels by lactate, but not by glucose, our data support both the notion that lactate is a preferred energy substrate over glucose for cortical neurons (Bouzier-Sore et al., 2003) and the astrocyte-neuron lactate shuttle hypothesis (Pellerin and Magistretti, 1994). Although the local cellular origin of lactate has been recently questioned (Lee et al., 2012;Diaz-Garcia et al., 2017), a growing number of evidence indicates that astrocytes are major central lactate producers (Almeida et al., 2001;Choi et al., 2012;Sotelo-Hitschfeld et al., 2015;Karagiannis et al., 2015;Le Douce J. et al., 2020;Jimenez-Blasco et al., 2020).

Glutamatergic synaptic transmission stimulates blood glucose uptake, astrocyte glycolysis, as well as lactate release (Pellerin and Magistretti, 1994;Voutsinos-Porche et al., 2003;Ruminot et al., 2011;Choi et al., 2012;Sotelo-Hitschfeld et al., 2015;Lerchundi et al., 2015) and diffusion through the astroglial gap junctional network (Rouach et al., 2008). This indicates that local and fast glutamatergic synaptic activity would be translated by astrocyte metabolism into a widespread and long-lasting extracellular lactate increase (Prichard et al., 1991;Hu and Wilson, 1997a), which could in turn enhance the firing of both excitatory and inhibitory neurons (Figure 7). Such a lactate surge would be spatially confined by the gap junctionnal connectivity of the astroglial network, which in layer IV represents an entire barrel (Houades et al., 2008).

This suggests that increased astrocytic lactate induced by whisker stimulation could enhance the activity of the cortical network and fine-tune upcoming sensory processing for several minutes, thereby favoring neuronal plasticity. Along this line, lactate derived from astrocyte glycogen supports both neuronal activity and long-term memory formation (Suzuki et al., 2011;Choi et al., 2012;Vezzoli et al., 2020). Similarly, cannabinoids, which notably alter neuronal processing and memory formation (Stella et al., 1997), hamper lactate production by astrocytes (Jimenez-Blasco et al., 2020).

Peripheral lactate produced by skeletal muscles during intense physical exercise and consumed by the active brain (Quistorff et al., 2008) could also have a similar but more global effect on neuronal processing and plasticity. Blood-borne lactate has been shown to promote learning and memory formation via brain-derived neurotrophic factor (El Hayek L. et al., 2019). It is worth noting that the production of this neurotrophin is altered in Kir6.2^-/-^ mice and impaired by a K_ATP_ channel opener (Fan et al., 2016), both conditions compromising the effect of lactate on spiking activity. Hence, the increase in astrocyte or systemic lactate could fine-tune neuronal processing and plasticity in a context-dependent manner and their coincidence could be potentially synergistic.

### Lactate-sensing compensatory mechanisms

Since excitatory neuronal activity increases extracellular lactate (Prichard et al., 1991;Hu and Wilson, 1997a) and lactate enhances neuronal activity, such a positive feedback loop (Figure 7) suggests that compensatory mechanisms might be recruited to prevent an overexcitation of neuronal activity by lactate supply. A metabolic negative feedback mechanism could involve the impairment of astrocyte metabolism and lactate release by endocannabinoids (Jimenez-Blasco et al., 2020) produced during intense neuronal activity (Stella et al., 1997).

Another possibility would consist in a blood flow decrease that would in turn reduce the delivery of blood glucose and subsequent local lactate production and release but also blood-borne lactate. Some GABAergic interneuron subtypes (Cauli et al., 2004;Uhlirova et al., 2016;Krawchuk et al., 2019), but also astrocytes (Girouard et al., 2010), can trigger vasoconstriction and blood flow decrease when their activity is increased. This could provide a negative hemodynamic feedback restricting spatially and temporally the increase of spiking activity by lactate.

PV-expressing and SOM-expressing interneurons exhibit higher mitochondrial content and apparent oxidative phosphorylation than pyramidal cells (Gulyas et al., 2006) suggesting that interneurons would more rapidly metabolize and sense lactate than pyramidal cells. These inhibitory GABAergic interneurons might therefore silence the cortical network, thereby providing a negative neuronal feedback loop. Active decrease in blood flow is associated with a decrease in neuronal activity (Shmuel et al., 2002;Shmuel et al., 2006;Devor et al., 2007). Vasoconstrictive GABAergic interneurons may underlie for both processes and could contribute to returning the system to a low lactate state.

## Conclusion

Our data indicate that lactate is both an energy substrate for cortical neurons and a signaling molecule enhancing their spiking activity. This suggests that a coordinated neurovascular and neurometabolic coupling would define a time window of an up state of lactate that, besides providing energy and maintenance to the cortical network, would fine-tune neuronal processing and favor, for example, memory formation (Suzuki et al., 2011;Kann et al., 2014;Galow et al., 2014;Jimenez-Blasco et al., 2020).

## Acknowledgments

This work was supported by grants from the Human Frontier Science Program (HFSP, RGY0070/2007, BC), the Agence Nationale pour la Recherche (ANR 2011 MALZ 003 01, BC). AK was supported by a Fondation pour la Recherche Médicale fellowship (FDT20100920106). We thank the animal facility of the IBPS (Paris, France).

## Author contributions

Conceptualization, T.G., J. Roeper and B.C.; Formal Analysis, J.N., B.C.; Investigation, A.K., T.G., A.L., F.P., J.P., H.G., R.H., B.C., Resources, B.L., J. Rossier, H.I., S.S., J. Roeper, B.C.; Writing –Original Draft, B.C.; Writing –Review & Editing, J.N., B.L.G., R.E., D.L., B.L., J.F.S., J. Roeper, B.C., Visualization, A.K., B.C.; Supervision, B.C.; Project Administration, B.C., Funding Acquisition, B.C.

## Materials and methods

### Key resources table

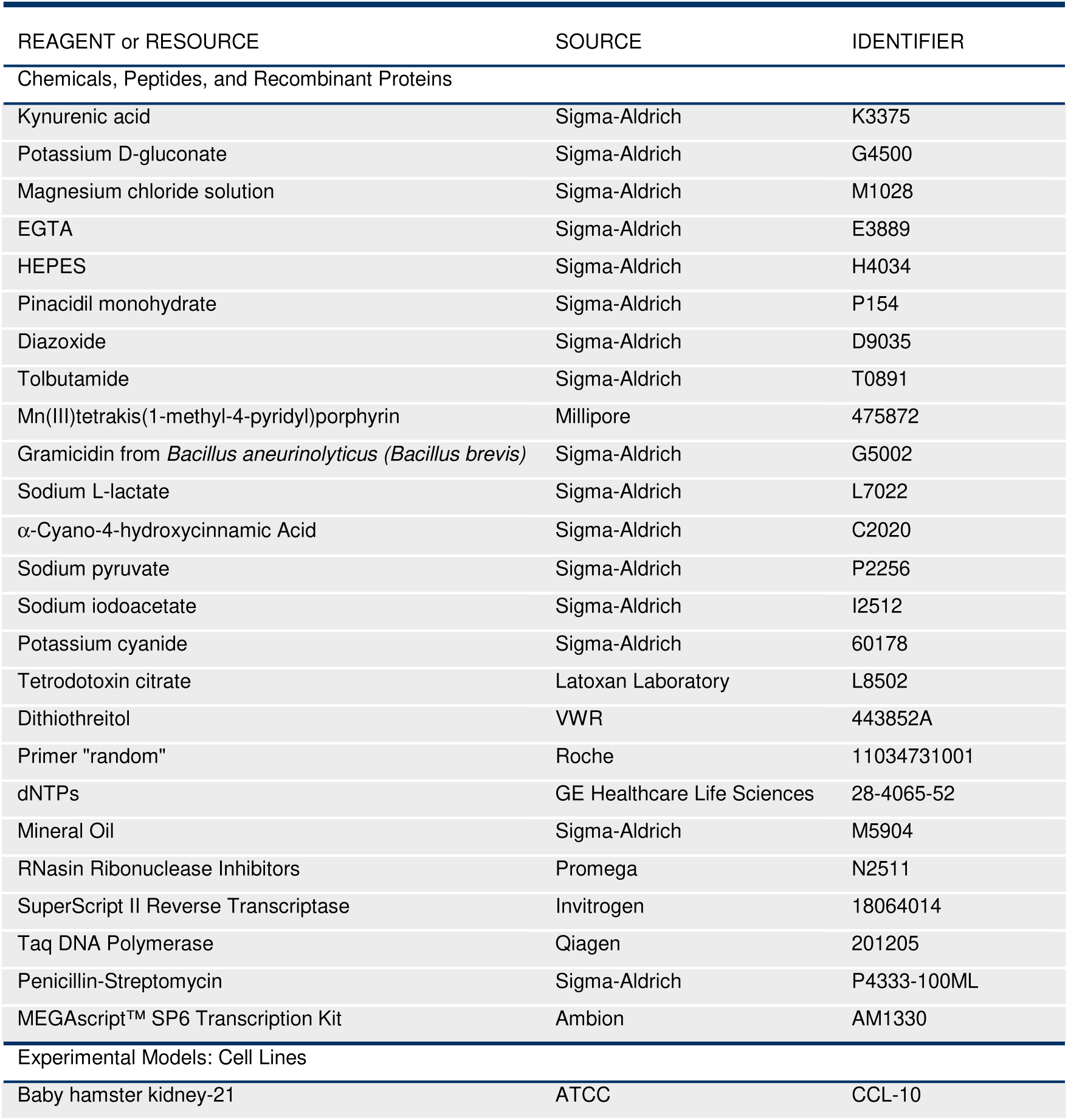
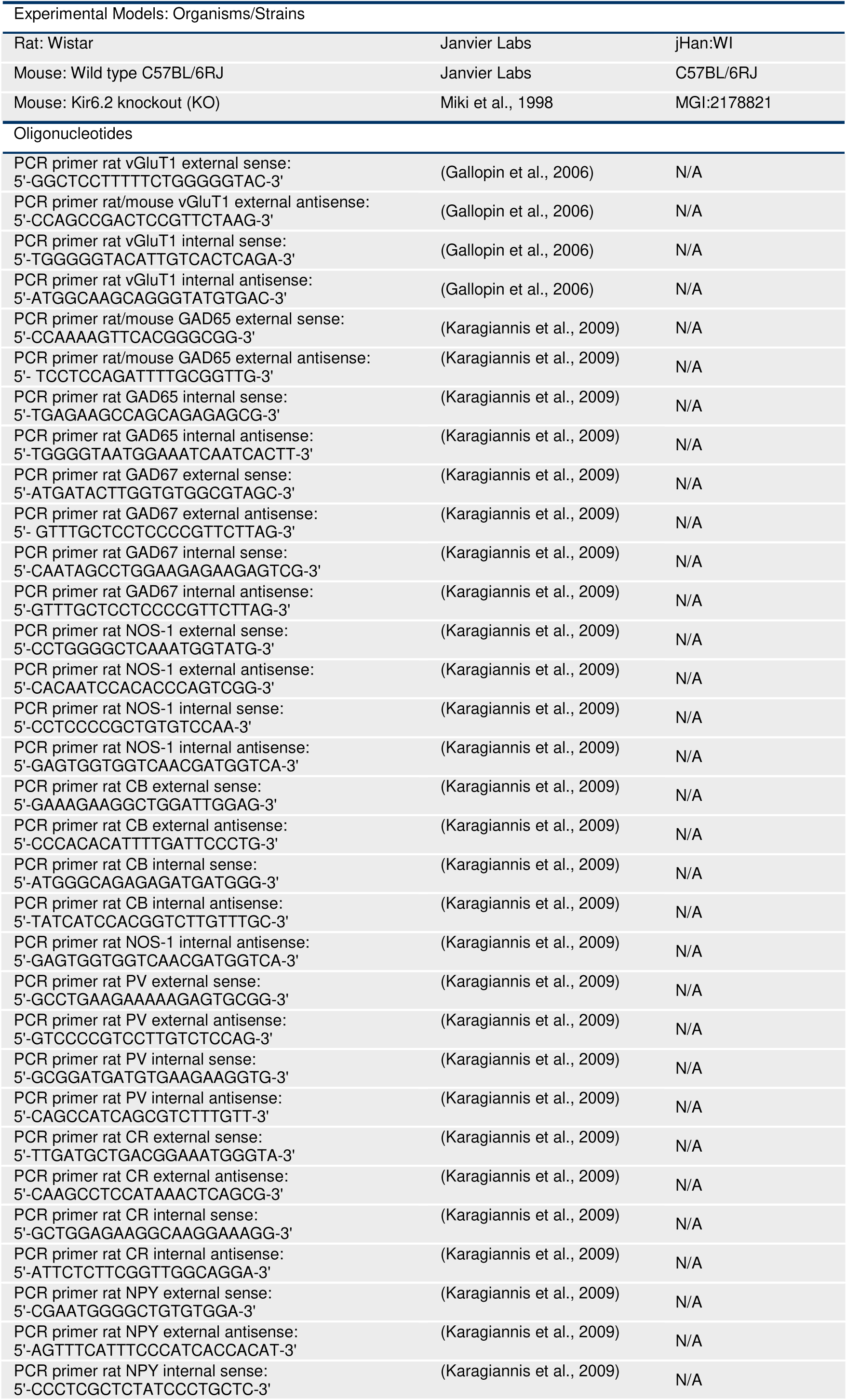
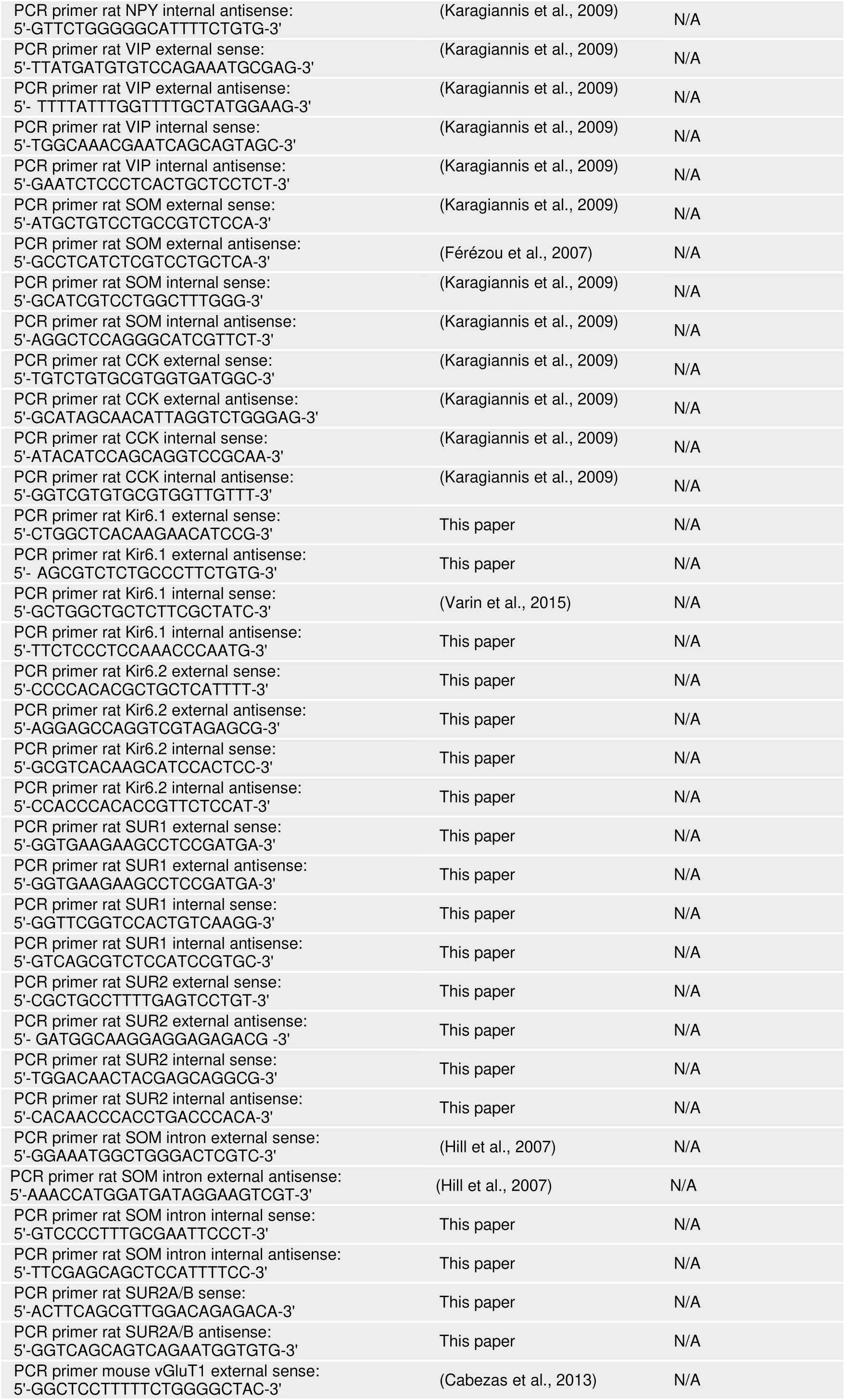
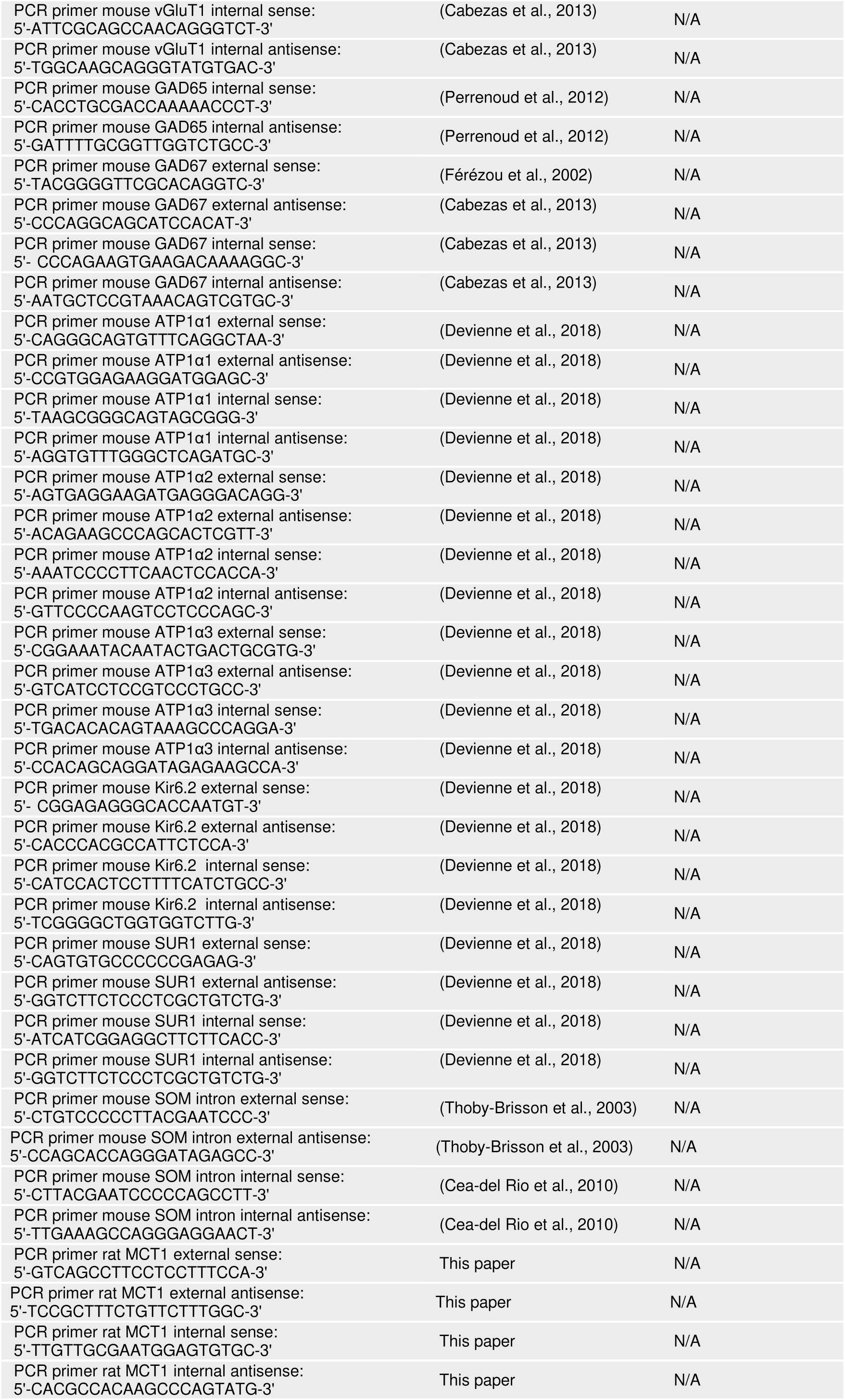
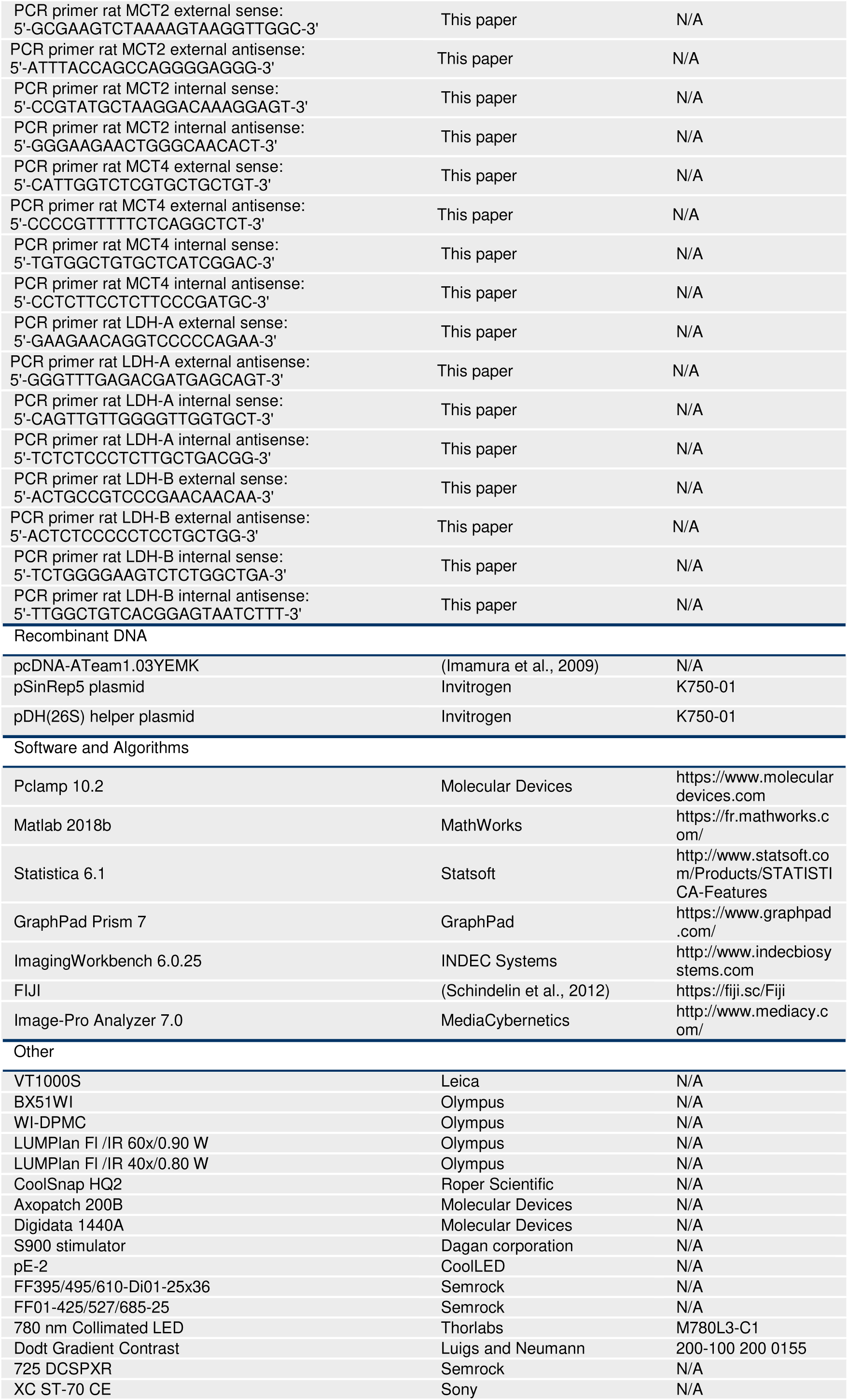
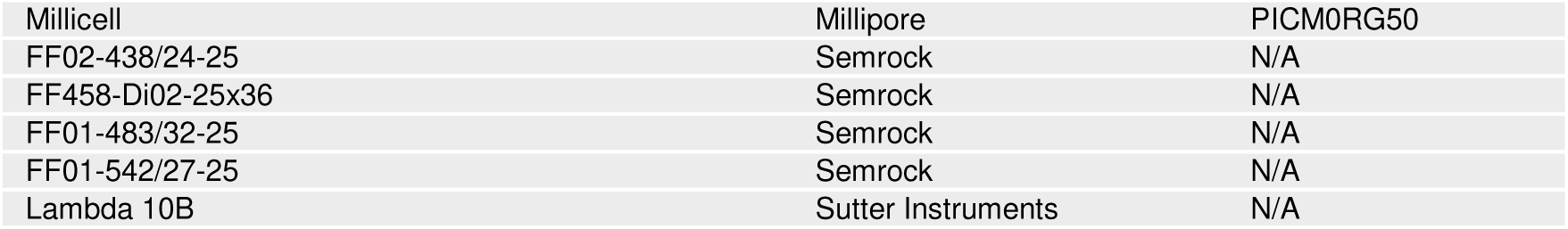

### Lead contact and materials availability

Further information and requests for resources and reagents should be directed to, and will be fulfilled by, the lead contact, B. Cauli.

### Experimental model and subject details

Wistar rats, C57BL/6RJ or Kir6.2 KO mice were used for all experiments in accordance with French regulations (Code Rural R214/87 to R214/130) and conformed to the ethical guidelines of both the directive 2010/63/EU of the European Parliament and of the Council and the French National Charter on the ethics of animal experimentation. A maximum of 3 rats or 5 mice were housed per cage and single animal housing was avoided. Male rats and mice of both genders were housed on a 12-hour light/dark cycle in a temperature-controlled (21–25°C) room and were given food and water *ad libitum.* Animals were used for experimentation at 13-24 days of age.

### Cortical slice preparation

Rats or mice were deeply anesthetized with isoflurane. After decapitation brains were quickly removed and placed into cold (∼4°C) oxygenated artificial cerebrospinal fluid (aCSF) containing (in mM): 126 NaCl, 2.5 KCl, 1.25 NaH_2_PO_4_, 2 CaCl_2_, 1 MgCl_2_, 26 NaHCO_3_, 10 glucose, 15 sucrose, and 1 kynurenic acid. Coronal slices (300 µm thick) containing the barrel cortex were cut with a vibratome (VT1000S, Leica) and allowed to recover at room temperature for at least 1h in aCSF saturated with O_2_/CO_2_ (95 %/5 %) as previously described (Karagiannis et al., 2009;Devienne et al., 2018).

### Whole-cell patch-clamp recording

Patch pipettes (4-6 MΩ) pulled from borosilicate glass were filled with 8 µl of RNAse free internal solution containing in (mM): 144 K-gluconate, 3 MgCl_2_, 0.5 EGTA, 10 HEPES, pH 7.2 (285/295 mOsm). Whole-cell recordings were performed at 25.3 ± 0.2°C using a patch-clamp amplifier (Axopatch 200B, Molecular Devices). Data were filtered at 5-10 kHz and digitized at 50 kHz using an acquisition board (Digidata 1440, Molecular Devices) attached to a personal computer running pCLAMP 10.2 software package (Molecular Devices). For ATP washout experiments neurons were recorded in voltage clamp mode using an ATP-free internal solution containing in (mM): 140 KCl, 20 NaCl, 2 MgCl_2_, 10 EGTA, 10 HEPES, pH 7.2.

### Cytoplasm harvesting and scRT-PCR

At the end of the whole-cell recording, lasting less than 15 min, the cytoplasmic content was aspirated in the recording pipette. The pipette’s content was expelled into a test tube and reverse transcription (RT) was performed in a final volume of 10 µl, as described previously (Lambolez et al., 1992). The scRT-PCR protocol was designed to probe simultaneously the expression of neuronal markers, K_ATP_ channels subunits or some key elements of energy metabolism. Two-steps amplification was performed essentially as described (Cauli et al., 1997;Devienne et al., 2018). Briefly, cDNAs present in the 10 µl reverse transcription reaction were first amplified simultaneously using all external primer pairs listed in the Key Ressources Table. Taq polymerase and 20 pmol of each primer were added to the buffer supplied by the manufacturer (final volume, 100 µl), and 20 cycles (94°C, 30 s; 60°C, 30 s; 72°C, 35 s) of PCR were run. Second rounds of PCR were performed using 1 µl of the first PCR product as a template. In this second round, each cDNA was amplified individually using its specific nested primer pair (Key Ressources Table) by performing 35 PCR cycles (as described above). 10 µl of each individual PCR product were run on a 2 % agarose gel stained with ethidium bromide using ФX174 digested by *Hae*III as a molecular weight marker.

### Perforated patch-clamp recording

Gramicidin stock solution (2 mg/ml, Sigma-Aldrich) was prepared in DMSO and diluted to 10-20 µg/ml (Zawar and Neumcke, 2000) in the RNAse free internal solution described above. The pipette tip was filled with gramicidin-free solution. Progress in perforation was evaluated by monitoring the capacitive transient currents elicited by −10 mV voltage pulses from a holding potential of −60 mV. In perforated patch configuration, a continuous current (52 ± 7 pA) was injected to induce the spiking of action potentials at stable firing rates of 4.1 ± 0.4 Hz obtained after an equilibration period of 3.6 ± 0.5 min. Membrane and access resistance were continuously monitored by applying −50 pA hyperpolarizing current pulses lasting 1 s every 10 s using an external stimulator (S900, Dagan) connected to the amplifier. Recordings were stopped when going into whole-cell configuration occurred, as evidenced by sudden increase of spike amplitude and decrease of access resistance.

### NADH imaging

Recordings were made in layer II-III of the rat somatosensory cortex. Wide-field fluorescent images were obtained using a double port upright microscope BX51WI, WI-DPMC, Olympus) with a 60x objective (LUMPlan Fl /IR 60x/0.90 W, Olympus) and a digital camera (CoolSnap HQ2, Roper Scientific) attached on the front port of the microscope. NADH autofluorescence was obtained by 365 nm excitation with a Light Emitting Device (LED, pE-2, CoolLED) using Imaging Workbench 6.0.25 software (INDEC Systems) and dichroic (FF395/495/610-Di01-25×36, Semrock) and emission filters (FF01-425/527/685-25, Semrock). Infrared Dodt gradient contrast images (IR-DGC, (Dodt and Zieglgansberger, 1998)) were obtained using a 780 nm collimated LED (M780L3-C1,Thorlabs) as a transmitted light source and DGC optics (Luigs and Neumann). Autofluorescence and IR-DGC images were collected every 10s by alternating the fluorescence and transmitted light sources. In parallel, infrared transmitted light images of slices were also continuously monitored on the back-port of the microscope using a customized beam splitter (725 DCSPXR, Semrock) and an analogic CCD camera (XC ST-70 CE, Sony). The focal plane was maintained constant on-line using infrared DGC images of cells as anatomical landmarks (Lacroix et al., 2015).

### Subcloning and viral production

The coding sequence of the ATP sensor ATeam1.03YEMK (Imamura et al., 2009) was subcloned into the viral vector pSinRep5. Sindbis virus was produced as previously described (Piquet et al., 2018). Recombinant pSinRep5 and helper plasmid pDH26S (Invitrogen) were transcribed in vitro into capped RNA using the Megascript SP6 kit (Ambion). Baby hamster kidney-21 cells (ATCC) were electroporated with sensor-containing RNA and helper RNA (2.10^7^ cells, 950 µF, 230 V) and incubated for 24 h at 37°C in 5% CO_2_ in Dulbecco’s modified Eagle Medium supplemented with 5% fetal calf serum before collecting cell supernatant containing the viruses. The virus titer (10^8^ infectious particles/ml) was determined after counting fluorescent baby hamster kidney cells infected using serial dilution of the stock virus.

### Brain slice viral transduction

Brain slices were placed onto a millicell membrane (Millipore) with culture medium (50% minimum essential medium, 50% Hank’s balanced salt sodium, 6.5 g/l glucose and 100 U/ml penicillin-streptomycin (Sigma-Aldrich) as previously described (Piquet et al., 2018). Infection was performed by adding ∼5 x 10^5^ particles per slice. Slices were incubated overnight at 35°C in 5% CO2. The next morning, brain slices were equilibrated for 1h in aCSF containing (in mM): 126 NaCl, 2.5 KCl, 1.25 NaH_2_PO_4_, 2 CaCl_2_, 1 MgCl_2_, 26 NaHCO_3_, 10 glucose, 15 sucrose. Slices were then placed into the recording chamber, heated at ∼30 °C and continuously perfused at 1-2 ml/min.

### FRET imaging

Recordings were made from visually identified pyramidal cells in layer II-III of the rat somatosensory cortex. Wide-field fluorescent images were obtained using a 40x objective and a digital camera attached on the front port of the microscope. The ATP sensor ATeam1.03YEMK was excited at 400 nm with a LED using Imaging Workbench 6.0.25 software and excitation (FF02-438/24-25, Semrock) and dichroic filters (FF458-Di02-25×36, Semrock). Double fluorescence images were collected every 15s by alternating the fluorescence emission filters for the CFP (FF01-483/32-25, Semrock) and the YFP (FF01-542/27-25, Semrock) using a filter wheel (Lambda 10B, Sutter Instruments). The focal plane was maintained constant on-line as described above.

### Pharmacological studies

Pinacidil (100 µM, Sigma-Aldrich); Diazoxide (300 µM, Sigma-Aldrich) and Tolbutamide (500 µM, Sigma-Aldrich), Mn(III)tetrakis(1-methyl-4-pyridyl)porphyrin (MnTMPyP, 25 µM, Millipore), α-cyano-4-hydroxycinnamate (4-CIN, 250 µM, Sigma-Aldrich); iodoacetic acid (IAA, 200 µM, Sigma-Aldrich) or KCN (1 mM, Sigma-Aldrich) was dissolved in aCSF from stock solutions of pinacidil (100 mM; NaOH 1M), diazoxide (300 mM; NaOH 1M), tolbutamide (500 mM; NaOH 1M), 4-CIN (250 mM; DMSO), IAA (200 mM, water) and KCN (1 M, water). Changes in extracellular glucose, lactate or pyruvate concentration were compensated by changes in sucrose concentration to maintain the osmolarity of the aCSF constant as previously described (Miki et al., 2001;Varin et al., 2015;Piquet et al., 2018) and pH was adjusted to 7.4.

### Quantification and statistical analysis

#### Analysis of somatic features

The laminar location determined by infrared videomicroscopy and recorded as 1-4 according to a location right within layers I, II/III or IV. For neurons located at the border of layers I-II/III and II/III-IV, the laminar location was represented by 1.5 and 3.5, respectively. Somatic features were measured from IR DGC of the recorded neurons. Briefly, the soma was manually delineated using Image-Pro Analyzer 7.0 software (MediaCybernetics) and length of major and minor axes, perimeter and area were extracted. The soma elongation was calculated as the ratio between major and minor axis. Roundness was calculated according to: 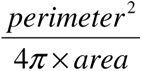; a value close to 1 is indicative of round somata.

### Analysis of electrophysiological properties

32 electrophysiological properties chosen to describe the electrophysiological diversity of cortical neurons (Ascoli et al., 2008) were determined using the I-clamp fast mode of the amplifier as previously described (Karagiannis et al., 2009). Membrane potential values were corrected for theoretical liquid junction potential (−15.6 mV). Resting membrane potential was measured just after passing in whole-cell configuration, and only cells with a resting membrane potential more negative than −55 mV were analyzed further. Membrane resistance (R_m_) and membrane time constant (τ_m_) were determined on responses to hyperpolarizing current pulses (duration, 800 ms) eliciting voltage shifts of 10-15 mV negative to rest (Kawaguchi, 1993;Kawaguchi, 1995). Time constant was determined by fitting this voltage response to a single exponential. Membrane capacitance (C_m_) was calculated according to C_m_=τ_m_/R_m_. Sag index was quantified as a relative decrease in membrane conductance according to (G_sag_-G_hyp_)/G_sag_ (Halabisky et al., 2006) where G_hyp_ and G_sag_ correspond to the whole-cell conductance when the sag was inactive and active, respectively. G_sag_ was measured as the slope of the linear portion of a current–voltage (I–V) plot, where V was determined at the end of 800 ms hyperpolarizing current pulses (−100 to 0 pA) and G_hyp_ as the slope of the linear portion of an I–V plot, where V was determined as the maximal negative potential during the 800 ms hyperpolarizing pulses. Rheobase was quantified as the minimal depolarizing current pulse intensity (800 ms duration pulses, 10 pA increments) generating at least one action potential. First spike latency (Gupta et al., 2000;Ascoli et al., 2008) was measured at rheobase as the time needed to elicit the first action potential. To describe different firing behaviors near threshold, spike frequency was measured near spike threshold on the first trace in which at least three spikes were triggered. Instantaneous discharge frequencies were measured and fitted to a straight line according to F_threshold_ = m_threshold_.t + F_min.,_ where m_threshold_ is the slope termed adaptation, t the time and F_min_, the minimal steady state frequency. Analysis of the action potentials waveforms was done on the first two spikes. Their amplitude (A1 and A2) was measured from threshold to the positive peak of the spike. Their duration (D1 and D2) was measured at half amplitude (Kawaguchi, 1993;Cauli et al., 1997). Their amplitude reduction and the duration increase were calculated according to (A1-A2)/A1 and (D2-D1)/D1, respectively (Cauli et al., 1997;Cauli et al., 2000). The amplitude and the latency of the fast and medium afterhyperpolarization (fAH and mAH) were measured for the first two action potentials as the difference between spike threshold and the negative peak of the AHs (Kawaguchi, 1993). The amplitude and latency of afterdepolarization (AD) following single spikes (Haj-Dahmane and Andrade, 1997) were measured as the difference between the negative peak of the fAH and the peak of the AD and between the spike threshold and the peak of the AD, respectively. When neurons did not exhibit mAH or AD, amplitude and latency were arbitrarily set to 0. A complex spike amplitude accommodation during a train of action potentials, consisting in a transient decrease of spikes amplitude, was measured as the difference between the peak of the smallest action potential and the peak of the following largest action potential (Cauli et al., 2000). Maximal firing rate was defined as the last trace before prominent reduction of action potentials amplitude indicative of a saturated discharge. To take into account the biphasic spike frequency adaptation (early and late) occurring at high firing rates (Cauli et al., 1997;Cauli et al., 2000;Gallopin et al., 2006), instantaneous firing frequency was fitted to a single exponential (Halabisky et al., 2006) with a sloping baseline, according to : *F_Saturation_* · = A_sat_ · e^-t/τsat^ + *t*.m_sat_ + F_max_, where A_sat_ corresponds to the amplitude of early frequency adaptation, τ_sat_ to the time constant of early adaptation, m_sat_ to the slope of late adaptation and F_max_ to the maximal steady state frequency.

### Unsupervised clustering

To classify neurons unsupervised clustering was performed using the laminar location of the soma, 10 molecular parameters (vGluT1, GAD65 and/or GAD67, NOS-1, CB, PV, CR, NPY, VIP, SOM and CCK) and the 32 electrophysiological parameters described above. Neurons positive for GAD65 and/or GAD67 were denoted as GAD positive and these mRNAs were considered as a single molecular variable as previously described (Gallopin et al., 2006). Parameters were standardized by centering and reducing all of the values. Cluster analysis was run on Statistica 6.1 software (Statsoft) using Ward’s method (Ward, 1963). The final number of clusters was established by hierarchically subdividing the clustering tree into higher order clusters as previously described (Karagiannis et al., 2009).

### Analysis of voltage clamp recordings

Whole-cell currents were measured from a holding potential of −70 mV and membrane resistances were determined by applying a voltage step to −60 mV of 100 ms every 5 s. The effects of K_ATP_ channel modulators were measured at the end of drug application by averaging, over a period of 1 minute, whole cell currents and changes in membrane resistance relative to control baseline prior to the application of drugs. Whole-cell K_ATP_ current and conductance were determined by subtracting current and conductance measured under K_ATP_ channel activator by their value measured under K_ATP_ channel blocker. The relative whole-cell K_ATP_ conductance was determined by dividing the whole-cell K_ATP_ conductance by the whole cell conductance measured under K_ATP_ channel activator. Whole-cell K_ATP_ current density was determined by dividing the whole-cell K_ATP_ current by the membrane capacitance. K_ATP_ current reversal potential was measured by subtracting I/V relationships obtained during voltage ramps from −60 to −130 mV determined under K_ATP_ channel activator and blocker, respectively.

During ATP washout experiments, whole-cell currents and I/V relationships were measured every 10 s at a holding potential of −50 mV and during voltage ramps from −40 to −120 mV, respectively. Washout currents were determined by subtracting the whole-cell currents measured at the beginning and the end of the whole cell-recording, respectively.

### Analysis of current clamp recordings

Every 10 s, membrane potential and mean firing rate were measured and membrane resistances were determined from voltage responses induced by −50 pA currents pulses lasting 1 s. K_ATP_ voltage response and changes in membrane resistance and firing rate were determined by subtracting their value measured under K_ATP_ channel activator by their value measured under K_ATP_ channel blocker. Neurons were considered as responsive to K_ATP_ channel modulators if the K_ATP_ channel activator induced both a hyperpolarization and a decrease in membrane resistance reversed by the K_ATP_ channel blocker.

### Analysis of perforated patch recordings

Mean firing frequency was measured every 10 s. Quantification of spiking activity was determined by averaging firing frequency over a period of 5 min preceding a change in extracellular aCSF composition. Firing frequencies were normalized by the averaged mean firing frequency measured under control condition.

### NADH imaging

Shading correction was applied off-line on the NADH autofluorescence images using the “Shading Corrector” plugin of FIJI software (Schindelin et al., 2012) and a blank field reference image. To compensate for potential x-y drifts all IR-DGC images were realigned off-line using the “StackReg” and “TurboReg” plugins (Thevenaz et al., 1998) of FIJI software and the same registration was applied to the corrected NAD(P)H autofluorescence images. To determine somatic regions of interest (ROIs) the soma was manually delineated on IR-DGC images. The mean NADH autofluorescence was measured at each time point using the same ROIs. Variations of fluorescence intensity were expressed as the ratio (F-F0)/F0 where F corresponds to the mean fluorescence intensity in the ROI at a given time point, and F0 corresponds to the mean fluorescence intensity in the same ROI during the 5 min control baseline prior to changes in aCSF composition. Effect of monocarboxylate superfusion or oxidative phosphorylation blockade was quantified by averaging the normalized ratio (R/R0) during the last five minutes of drug application.

### FRET imaging

All images were realigned off-line as described above using the YFP images as the reference for registration. Fluorescence ratios were calculated by dividing the registered YFP images by the registered CFP images using FIJI. The somatic ROIs were manually delineated on the YFP images as described above. The mean ratio was measured at each time point using the same ROIs. Variations of fluorescence ratio were expressed as the ratio (R-R0)/R0 where R corresponds to the fluorescence ratio in the ROI at a given time point, and R0 corresponds to the mean fluorescence ratio in the same ROI during the 10 min control baseline prior to drug application. Effect of glycolysis or oxidative phosphorylation blockade was quantified by averaging the normalized ratio during the last five minutes of drug application.

### Statistical analysis

Statistical analyses were performed with Statistica 6.1 and GraphPad Prism 7. All values are expressed as means ± s.e.m. Normality of distributions and equality of variances were assessed using the Shapiro–Wilk test and the Fisher F-test, respectively. Parametric tests were only used if these criteria were met. Holm-Bonferroni correction was used for multiple comparisons and p-values are given as uncorrected. Statistical significance on all figures uses the following convention of corrected p-values: * p <0.05, ** p < 0.01, *** p <0.001.

Statistical significance of morphological and electrophysiological properties of neurons was determined using the Mann-Withney U test. Comparison of the occurrence of expressed genes and of responsiveness of K_ATP_ channel modulators between different cell types was determined using Fisher’s exact test. Statistical significance of the effects of K_ATP_ channel modulators was determined using the Friedman and post hoc Dunn’s tests. Significance of the effect of the ROS scavenger was determined using one-tailed unpaired student t-test. Comparison of K_ATP_ channel properties was determined using Mann-Withney U, Student-t, or Kruskal-Wallis H tests. Comparison of responses between Kir6.2^+/+^ and Kir6.2^-/-^ neurons was determined using Mann-Withney U test. Statistical significance of the effects of energy substrates and drug applications on evoked firing in perforated patch recordings was determined using Friedman and Dunn’s tests. Comparison of the effects of monocarboxylates and cyanide on NADH fluorescence was determined using Mann-Withney U test. Statistical significance of the effects of metabolic inhibitors on intracellular ATP was determined using Friedman and Dunn’s tests.

---

**Figure S1.**
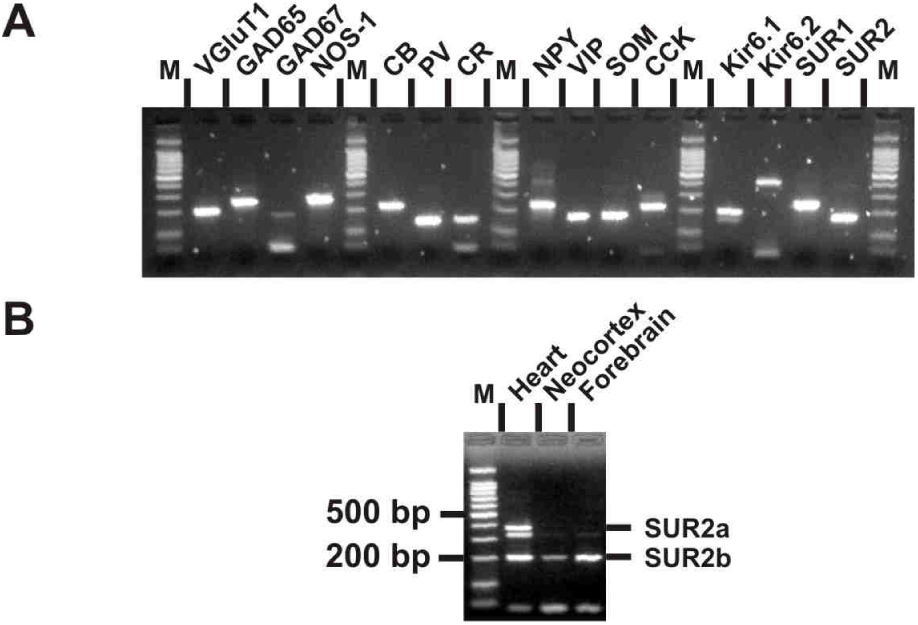
Molecular expression of K_ATP_ channels. (A) RT-PCR products generated from 500 pg of total cortical RNAs. M: 100 bp ladder molecular weight marker. (B) SUR2 splice variants-specific RT-PCR analysis of 1 ng total RNAs from rat heart, neocortex and forebrain.

**Figure S2.**
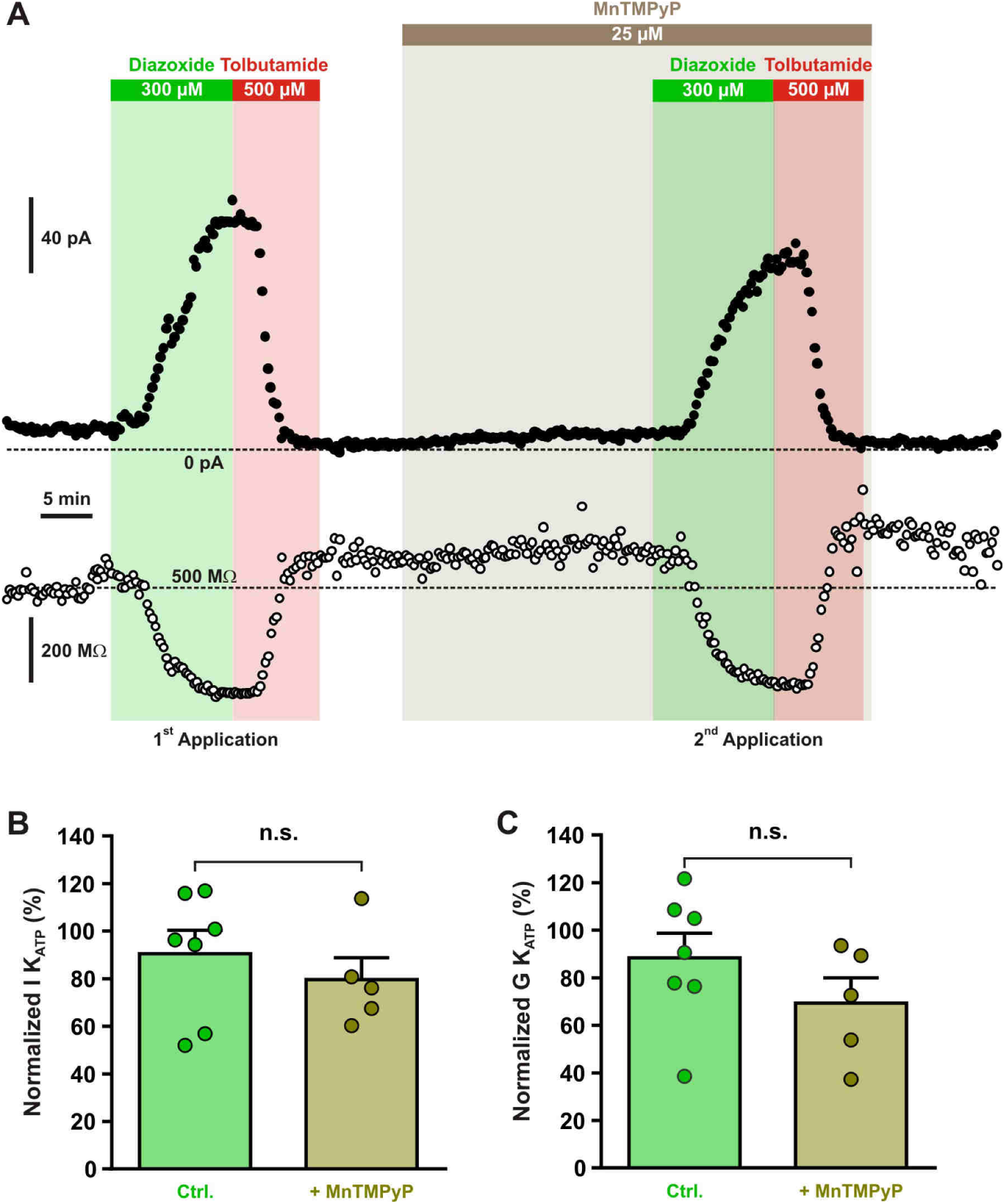
Diazoxide-induced current is independent of ROS production. (A) Representative stationary currents at −60 mV (filled circles) and membrane resistance (open circles) changes induced by diazoxide and tolbutamide under control condition and in presence of the superoxide dismutase and catalase mimetic, MnTMPyP. The colored bars and shaded zones indicate the duration of application. (B-C) Histograms summarizing the relative K_ATP_ currents (B) and relative whole-cell K_ATP_ conductance (C) evoked by two consecutive diazoxide and tolbutamide applications in control condition (Ctrl.) and after the presence of MnTMPyP. Data are normalized by the data measured during first application, expressed as mean ± s.e.m., and the individual data points are depicted. n.s. not statistically significant.

**Figure S3.**
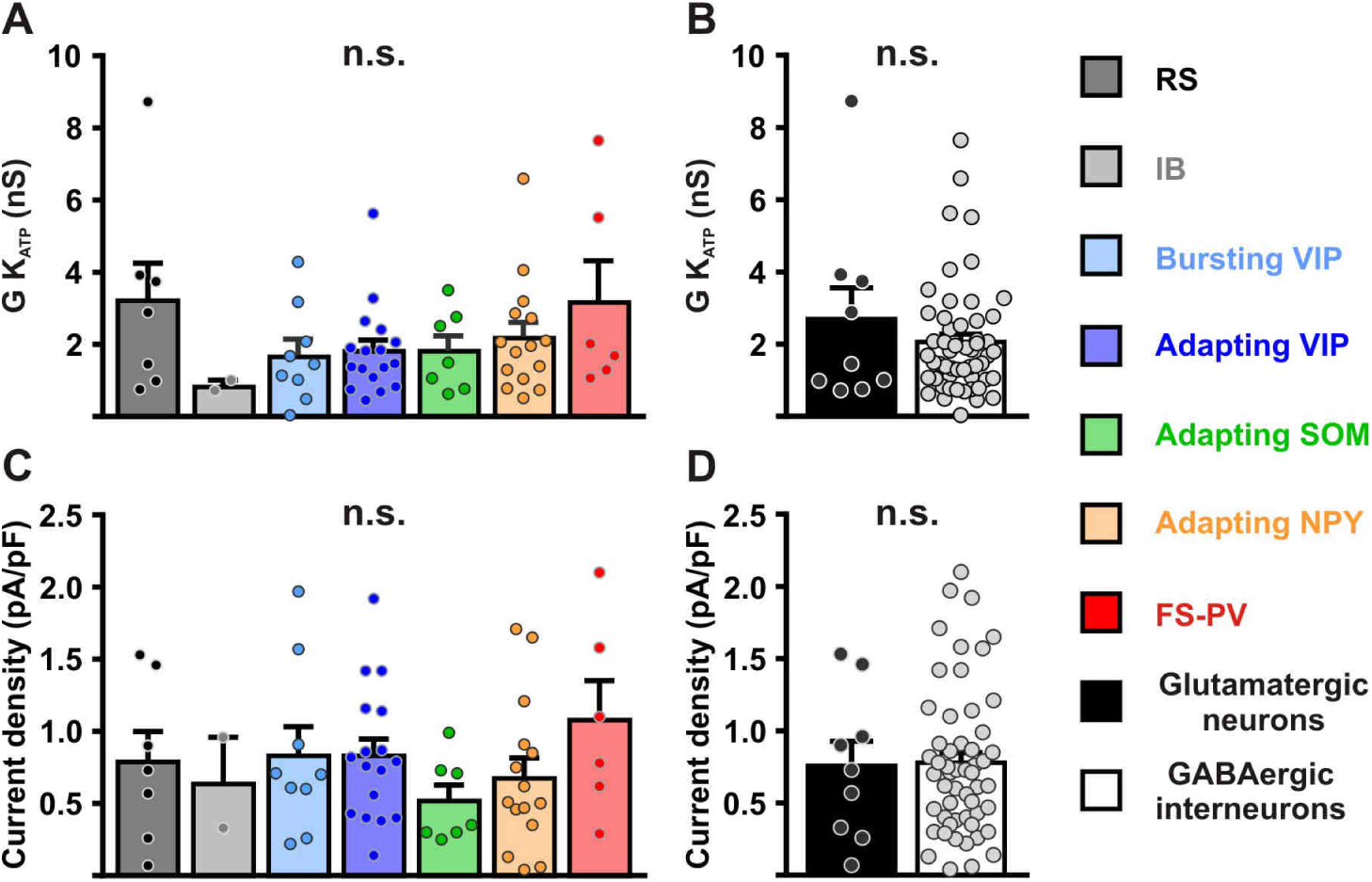
Characterization of K_ATP_ channels in different cortical neurons. (A-D) Histograms summarizing the whole-cell K_ATP_ conductance (A, B) and K_ATP_ current density (C, D) and K_ATP_ current reversal potential in identified neuronal subtypes (A,C) or between glutamatergic and GABAergic neurons (B,D). Data are expressed as mean ± s.e.m., and the individual data points are depicted. n.s. not statistically significant.

**Figure S4.**
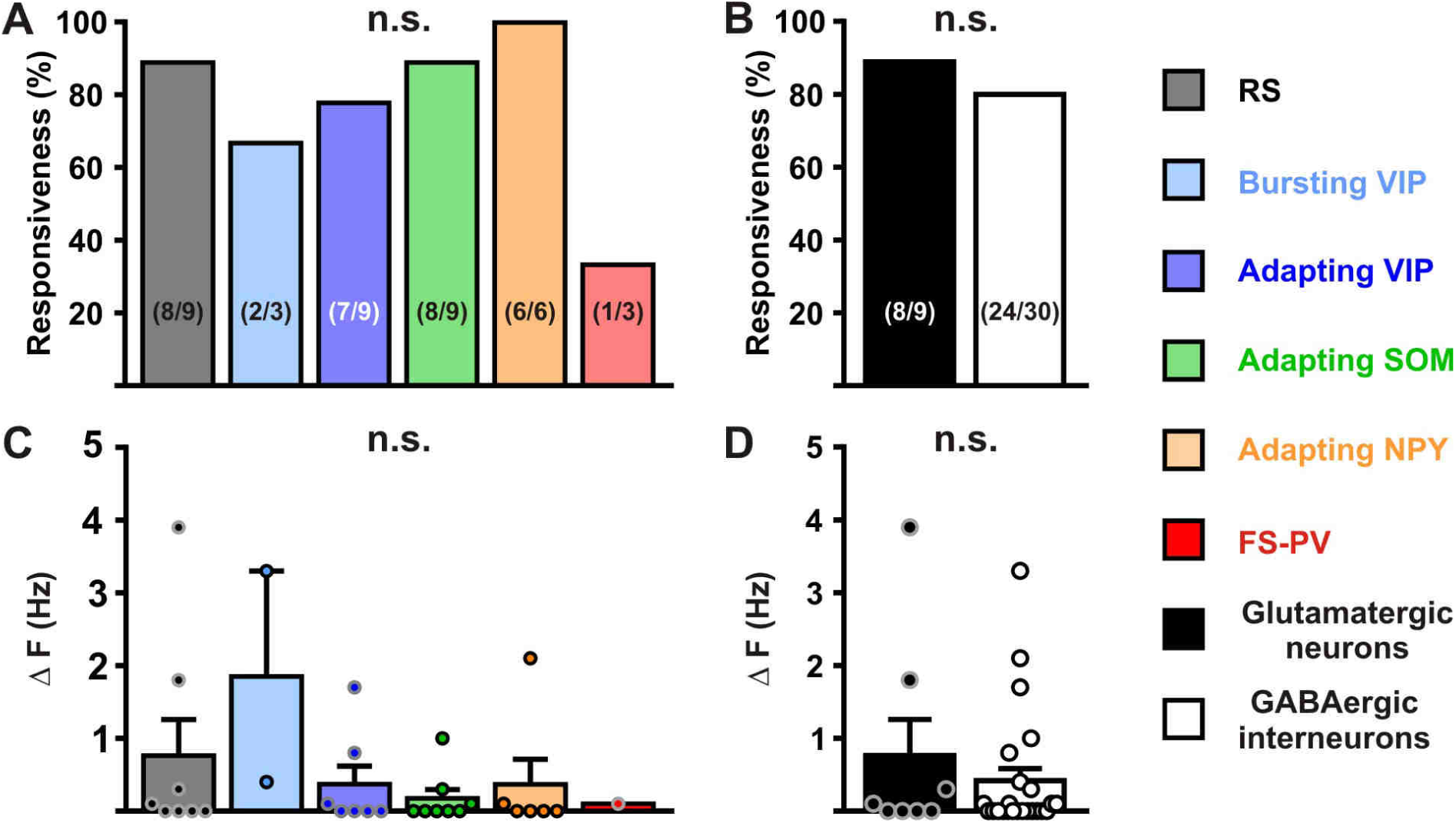
Modulation of neuronal activity in different cortical neurons by K_ATP_ channels. (A-D) Histograms summarizing the proportion of responsive neurons (A, K^2^_(5)_=7.3125, p=0.1984, and B, p=0.9999) and modulation firing rate (C, H_(5,32)_=5.69107, p=0.337, and D, U_(8,24)_=87.5, p=0.6994) by K_ATP_ channels in neuronal subtypes (A,C) and groups (B,D). The numbers in brackets indicate the number of responsive cells and analyzed cells, respectively. Data are expressed as mean ± s.e.m., and the individual data points are depicted. n.s. not statistically significant.

**Figure S5.**
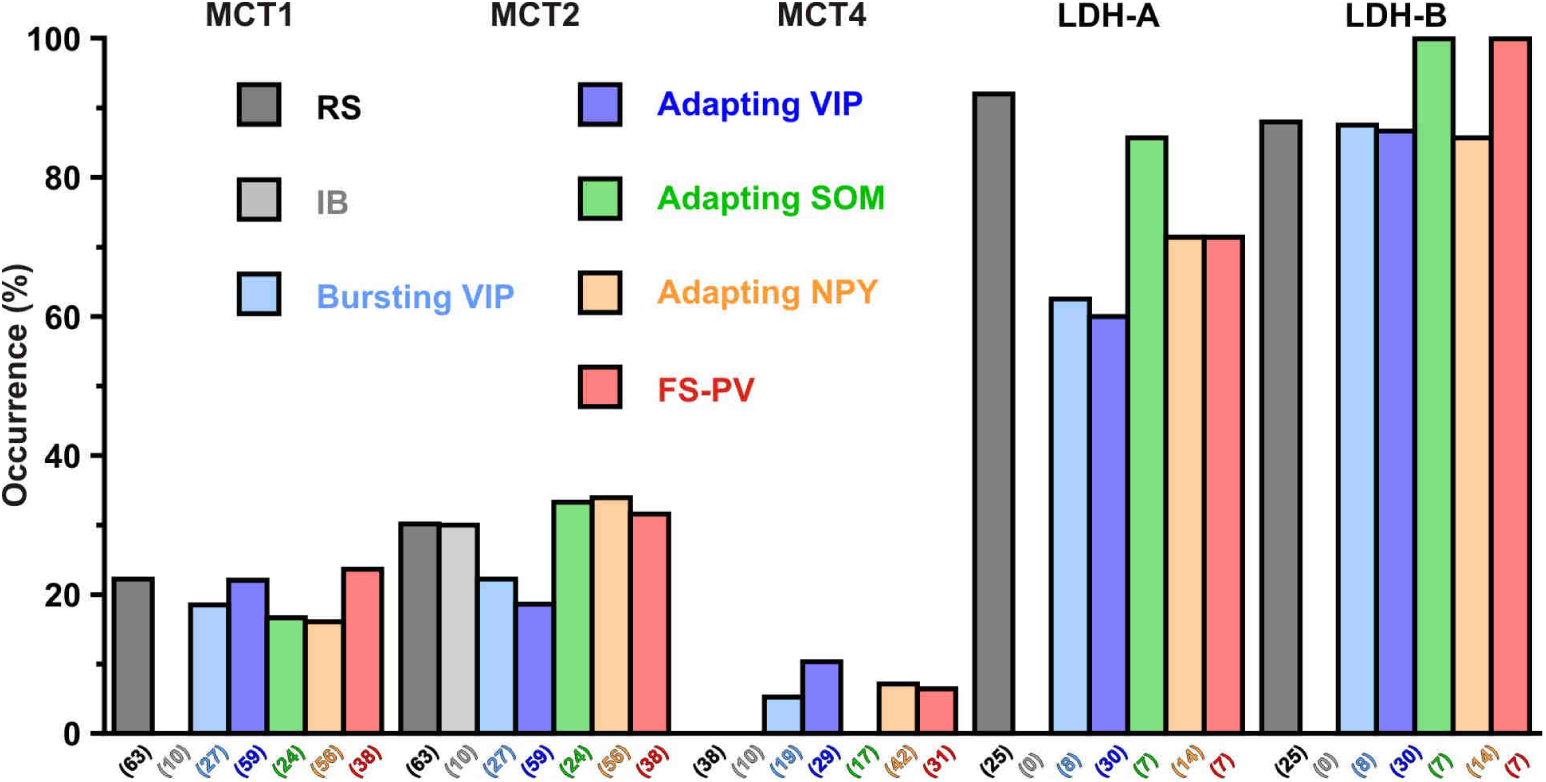
Expression profile of monocarboxylate transporters and lactate dehydrogenase subunits in different cortical neuronal types. Histograms summarizing the expression profile of the monocarboxylate transporters MCT1, 2 and 4 and LDH A and B lactate dehydrogenase subunits in different neuronal subtypes. The numbers in brackets indicate the number of analyzed cells.

**Figure S6.**
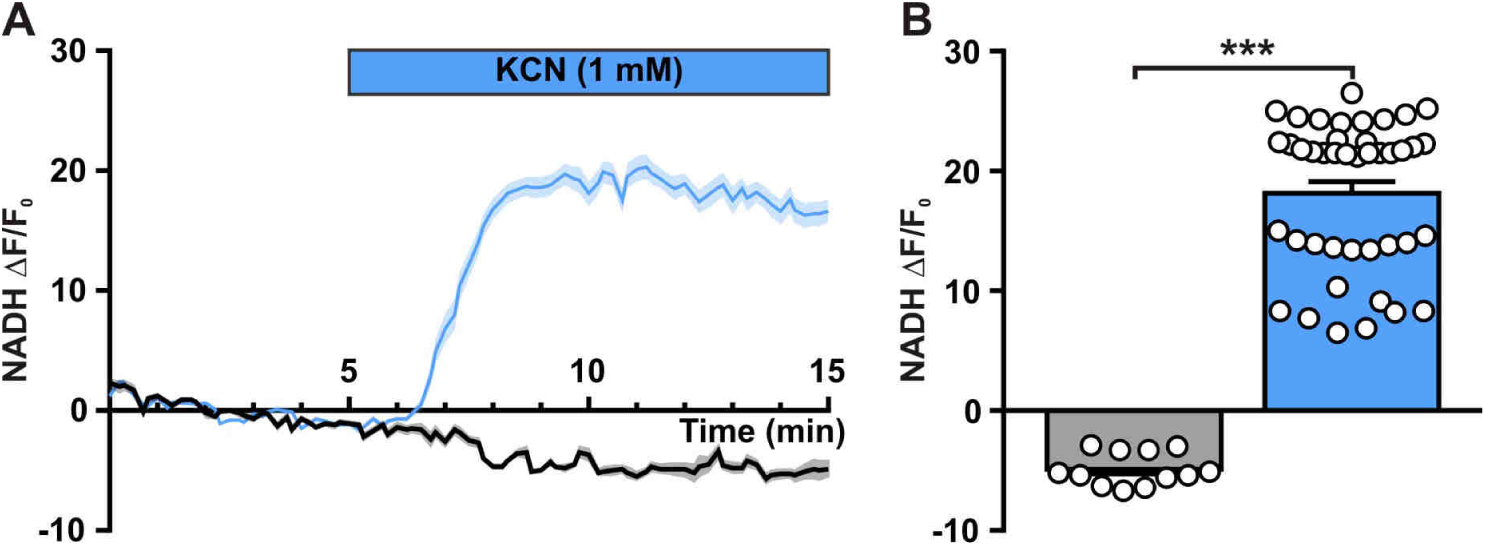
Neuronal NADH autofluorescence increase by blockade of oxidative phosphorylation. (A) Mean relative changes in NADH autofluorescence in control condition (grey) and in response to 1 mM KCN (blue). The colored bar indicates the duration of KCN applications. Data are expressed as mean ± s.e.m. (B) Histograms summarizing the mean relative changes in NADH autofluorescence measured during the last 5 minutes of 1 mM KCN application (blue) and corresponding time in control condition (grey). Data are expressed as mean ± s.e.m., and the individual data points are depicted.

## References

Aguilar-Bryan, L., Nichols, C.G., Wechsler, S.W., Clement, J.P., Boyd, A.E., III, Gonzalez, G., Herrera-Sosa, H., Nguy, K., Bryan, J., and Nelson, D.A. (1995). Cloning of the beta cell high-affinity sulfonylurea receptor: a regulator of insulin secretion. Science 268, 423–426.

Ahmed, K., Tunaru, S., Tang, C., Muller, M., Gille, A., Sassmann, A., Hanson, J., and Offermanns, S. (2010). An autocrine lactate loop mediates insulin-dependent inhibition of lipolysis through GPR81. Cell Metab 11, 311–319.

Ainscow, E.K., Mirshamsi, S., Tang, T., Ashford, M.L., and Rutter, G.A. (2002). Dynamic imaging of free cytosolic ATP concentration during fuel sensing by rat hypothalamic neurones: evidence for ATP-independent control of ATP-sensitive K(+) channels. J. Physiol 544, 429–445.

Almeida, A., Almeida, J., Bolanos, J.P., and Moncada, S. (2001). Different responses of astrocytes and neurons to nitric oxide: the role of glycolytically generated ATP in astrocyte protection. Proc. Natl. Acad. Sci. U. S. A 98, 15294–15299.

Ammala, C., Moorhouse, A., and Ashcroft, F.M. (1996). The sulphonylurea receptor confers diazoxide sensitivity on the inwardly rectifying K+ channel Kir6.1 expressed in human embryonic kidney cells. J. Physiol 494 (Pt 3), 709–714.

Ascoli, G.A., Alonso-Nanclares, L., Anderson, S.A., Barrionuevo, G., avides-Piccione, R., Burkhalter, A., Buzsaki, G., Cauli, B., DeFelipe, J., Fairen, A., Feldmeyer, D., Fishell, G., Fregnac, Y., Freund, T.F., Gardner, D., Gardner, E.P., Goldberg, J.H., Helmstaedter, M., Hestrin, S., Karube, F., Kisvarday, Z.F., Lambolez, B., Lewis, D.A., Marin, O., Markram, H., Munoz, A., Packer, A., Petersen, C.C., Rockland, K.S., Rossier, J., Rudy, B., Somogyi, P., Staiger, J.F., Tamas, G., Thomson, A.M., Toledo-Rodriguez, M., Wang, Y., West, D.C., and Yuste, R. (2008). Petilla terminology: nomenclature of features of GABAergic interneurons of the cerebral cortex. Nat. Rev. Neurosci. 9, 557–568.

Ashford, M.L., Sturgess, N.C., Trout, N.J., Gardner, N.J., and Hales, C.N. (1988). Adenosine-5’-triphosphate-sensitive ion channels in neonatal rat cultured central neurones. Pflugers Arch. 412, 297–304.

Attwell, D., and Laughlin, S.B. (2001). An energy budget for signaling in the grey matter of the brain. J. Cereb. Blood Flow Metab 21, 1133–1145.

Aziz, Q., Li, Y., Anderson, N., Ojake, L., Tsisanova, E., and Tinker, A. (2017). Molecular and functional characterization of the endothelial ATP-sensitive potassium channel. J. Biol. Chem. 292, 17587–17597.

Babenko, A.P., Aguilar-Bryan, L., and Bryan, J. (1998). A view of sur/KIR6.X, KATP channels. Annu. Rev. Physiol 60, 667–687.

Bittar, P.G., Charnay, Y., Pellerin, L., Bouras, C., and Magistretti, P.J. (1996). Selective distribution of lactate dehydrogenase isoenzymes in neurons and astrocytes of human brain. J. Cereb. Blood Flow Metab 16, 1079–1089.

Bondjers, C., He, L., Takemoto, M., Norlin, J., Asker, N., Hellstrom, M., Lindahl, P., and Betsholtz, C. (2006). Microarray analysis of blood microvessels from PDGF-B and PDGF-Rbeta mutant mice identifies novel markers for brain pericytes. FASEB J. 20, 1703–1705.

Bouzier-Sore, A.K., Voisin, P., Bouchaud, V., Bezancon, E., Franconi, J.M., and Pellerin, L. (2006). Competition between glucose and lactate as oxidative energy substrates in both neurons and astrocytes: a comparative NMR study. Eur. J. Neurosci. 24, 1687–1694.

Bouzier-Sore, A.K., Voisin, P., Canioni, P., Magistretti, P.J., and Pellerin, L. (2003). Lactate is a preferential oxidative energy substrate over glucose for neurons in culture. J. Cereb. Blood Flow Metab 23, 1298–1306.

Bozzo, L., Puyal, J., and Chatton, J.Y. (2013). Lactate modulates the activity of primary cortical neurons through a receptor-mediated pathway. PLoS. ONE. 8, e71721.

Broer, S., Broer, A., Schneider, H.P., Stegen, C., Halestrap, A.P., and Deitmer, J.W. (1999). Characterization of the high-affinity monocarboxylate transporter MCT2 in Xenopus laevis oocytes. Biochem. J. 341 (Pt 3), 529–535.

Broer, S., Schneider, H.P., Broer, A., Rahman, B., Hamprecht, B., and Deitmer, J.W. (1998). Characterization of the monocarboxylate transporter 1 expressed in Xenopus laevis oocytes by changes in cytosolic pH. Biochem. J. 333 (Pt 1), 167–174.

Cabezas, C., Irinopoulou, T., Cauli, B., and Poncer, J.C. (2013). Molecular and functional characterization of GAD67-expressing, newborn granule cells in mouse dentate gyrus. Front Neural Circuits. 7, 60.

Cahoy, J.D., Emery, B., Kaushal, A., Foo, L.C., Zamanian, J.L., Christopherson, K.S., Xing, Y., Lubischer, J.L., Krieg, P.A., Krupenko, S.A., Thompson, W.J., and Barres, B.A. (2008). A transcriptome database for astrocytes, neurons, and oligodendrocytes: a new resource for understanding brain development and function. J. Neurosci. 28, 264–278.

Cao, R., Higashikubo, B.T., Cardin, J., Knoblich, U., Ramos, R., Nelson, M.T., Moore, C.I., and Brumberg, J.C. (2009). Pinacidil induces vascular dilation and hyperemia in vivo and does not impact biophysical properties of neurons and astrocytes in vitro. Cleve. Clin. J. Med. 76 Suppl 2, S80–S85.

Cauli, B., Audinat, E., Lambolez, B., Angulo, M.C., Ropert, N., Tsuzuki, K., Hestrin, S., and Rossier, J. (1997). Molecular and physiological diversity of cortical nonpyramidal cells. J. Neurosci. 17, 3894–3906.

Cauli, B., Porter, J.T., Tsuzuki, K., Lambolez, B., Rossier, J., Quenet, B., and Audinat, E. (2000). Classification of fusiform neocortical interneurons based on unsupervised clustering. Proc. Natl. Acad. Sci. U. S. A 97, 6144–6149.

Cauli, B., Tong, X.K., Rancillac, A., Serluca, N., Lambolez, B., Rossier, J., and Hamel, E. (2004). Cortical GABA interneurons in neurovascular coupling: relays for subcortical vasoactive pathways. J. Neurosci. 24, 8940–8949.

Cea-del Rio, C.A., Lawrence, J.J., Tricoire, L., Erdelyi, F., Szabo, G., and McBain, C.J. (2010). M3 muscarinic acetylcholine receptor expression confers differential cholinergic modulation to neurochemically distinct hippocampal basket cell subtypes. J. Neurosci. 30, 6011–6024.

Chance, B., Cohen, P., Jobsis, F., and Schoener, B. (1962). Intracellular oxidation-reduction states in vivo. Science 137, 499–509.

Choi, H.B., Gordon, G.R., Zhou, N., Tai, C., Rungta, R.L., Martinez, J., Milner, T.A., Ryu, J.K., McLarnon, J.G., Tresguerres, M., Levin, L.R., Buck, J., and MacVicar, B.A. (2012). Metabolic Communication between Astrocytes and Neurons via Bicarbonate-Responsive Soluble Adenylyl Cyclase. Neuron 75, 1094–1104.

Chuquet, J., Quilichini, P., Nimchinsky, E.A., and Buzsaki, G. (2010). Predominant Enhancement of Glucose Uptake in Astrocytes versus Neurons during Activation of the Somatosensory Cortex. J. Neurosci. 30, 15298–15303.

Chutkow, W.A., Simon, M.C., Le Beau, M.M., and Burant, C.F. (1996). Cloning, tissue expression, and chromosomal localization of SUR2, the putative drug-binding subunit of cardiac, skeletal muscle, and vascular KATP channels. Diabetes 45, 1439–1445.

Clarke, D.D., and Sokoloff, L. (1999). Circulation and Energy Metabolism of the Brain. In Basic Neurochemistry: Molecular, Cellular and Medical Aspects., G.J. Siegel, ed. (Philadelphia: Lippincott Williams & Wilkins), pp. 637–669.

Cunningham, M.O., Pervouchine, D.D., Racca, C., Kopell, N.J., Davies, C.H., Jones, R.S., Traub, R.D., and Whittington, M.A. (2006). Neuronal metabolism governs cortical network response state. Proc. Natl. Acad. Sci. U. S. A 103, 5597–5601.

D’Agostino, D.P., Putnam, R.W., and Dean, J.B. (2007). Superoxide ({middle dot}O2-) production in CA1 neurons of rat hippocampal slices exposed to graded levels of oxygen. J. Neurophysiol.

de Castro Abrantes H., Briquet, M., Schmuziger, C., Restivo, L., Puyal, J., Rosenberg, N., Rocher, A.B., Offermanns, S., and Chatton, J.Y. (2019). The lactate receptor HCAR1 modulates neuronal network activity through the activation of Galpha and Gbeta subunits. J. Neurosci.

Devienne, G., Le Gac, B., Piquet, J., and Cauli, B. (2018). Single Cell Multiplex Reverse Transcription Polymerase Chain Reaction After Patch-clamp. J. Vis. Exp.

Devor, A., Hillman, E.M., Tian, P., Waeber, C., Teng, I.C., Ruvinskaya, L., Shalinsky, M.H., Zhu, H., Haslinger, R.H., Narayanan, S.N., Ulbert, I., Dunn, A.K., Lo, E.H., Rosen, B.R., Dale, A.M., Kleinfeld, D., and Boas, D.A. (2008). Stimulus-induced changes in blood flow and 2-deoxyglucose uptake dissociate in ipsilateral somatosensory cortex. J. Neurosci. 28, 14347–14357.

Devor, A., Tian, P., Nishimura, N., Teng, I.C., Hillman, E.M., Narayanan, S.N., Ulbert, I., Boas, D.A., Kleinfeld, D., and Dale, A.M. (2007). Suppressed neuronal activity and concurrent arteriolar vasoconstriction may explain negative blood oxygenation level-dependent signal. J. Neurosci. 27, 4452–4459.

Dhar-Chowdhury, P., Harrell, M.D., Han, S.Y., Jankowska, D., Parachuru, L., Morrissey, A., Srivastava, S., Liu, W., Malester, B., Yoshida, H., and Coetzee, W.A. (2005). The glycolytic enzymes, glyceraldehyde-3-phosphate dehydrogenase, triose-phosphate isomerase, and pyruvate kinase are components of the K(ATP) channel macromolecular complex and regulate its function. J. Biol. Chem. 280, 38464–38470.

Diaz-Garcia, C.M., Lahmann, C., Martinez-Francois, J.R., Li, B., Koveal, D., Nathwani, N., Rahman, M., Keller, J.P., Marvin, J.S., Looger, L.L., and Yellen, G. (2019). Quantitative in vivo imaging of neuronal glucose concentrations with a genetically encoded fluorescence lifetime sensor. J. Neurosci. Res.

Diaz-Garcia, C.M., Mongeon, R., Lahmann, C., Koveal, D., Zucker, H., and Yellen, G. (2017). Neuronal Stimulation Triggers Neuronal Glycolysis and Not Lactate Uptake. Cell Metab 26, 361–374.

Dodt, H.U., and Zieglgansberger, W. (1998). Visualization of neuronal form and function in brain slices by infrared videomicroscopy. Histochem. J. 30, 141–152.

Drose, S., Brandt, U., and Hanley, P.J. (2006). K+-independent actions of diazoxide question the role of inner membrane KATP channels in mitochondrial cytoprotective signaling. J. Biol. Chem. 281, 23733–23739.

Dufer, M., Krippeit-Drews, P., Buntinas, L., Siemen, D., and Drews, G. (2002). Methyl pyruvate stimulates pancreatic beta-cells by a direct effect on KATP channels, and not as a mitochondrial substrate. Biochem. J. 368, 817–825.

Dunn-Meynell, A.A., Rawson, N.E., and Levin, B.E. (1998). Distribution and phenotype of neurons containing the ATP-sensitive K+ channel in rat brain. Brain Res. 814, 41–54.

El Hayek L., Khalifeh, M., Zibara, V., Abi, A.R., Emmanuel, N., Karnib, N., El-Ghandour, R., Nasrallah, P., Bilen, M., Ibrahim, P., Younes, J., Abou, H.E., Barmo, N., Jabre, V., Stephan, J.S., and Sleiman, S.F. (2019). Lactate Mediates the Effects of Exercise on Learning and Memory through SIRT1-Dependent Activation of Hippocampal Brain-Derived Neurotrophic Factor (BDNF). J. Neurosci. 39, 2369–2382.

Fan, Y., Kong, H., Ye, X., Ding, J., and Hu, G. (2016). ATP-sensitive potassium channels: uncovering novel targets for treating depression. Brain Struct. Funct. 221, 3111–3122.

Férézou, I., Cauli, B., Hill, E.L., Rossier, J., Hamel, E., and Lambolez, B. (2002). 5-HT3 receptors mediate serotonergic fast synaptic excitation of neocortical vasoactive intestinal peptide/cholecystokinin interneurons. J. Neurosci. 22, 7389–7397.

Férézou, I., Hill, E.L., Cauli, B., Gibelin, N., Kaneko, T., Rossier, J., and Lambolez, B. (2007). Extensive overlap of mu-opioid and nicotinic sensitivity in cortical interneurons. Cereb. Cortex 17, 1948–1957.

Galeffi, F., Foster, K.A., Sadgrove, M.P., Beaver, C.J., and Turner, D.A. (2007). Lactate uptake contributes to the NAD(P)H biphasic response and tissue oxygen response during synaptic stimulation in area CA1 of rat hippocampal slices. J. Neurochem. 103, 2449–2461.

Gallopin, T., Geoffroy, H., Rossier, J., and Lambolez, B. (2006). Cortical sources of CRF, NKB, and CCK and their effects on pyramidal cells in the neocortex. Cereb. Cortex 16, 1440–1452.

Galow, L.V., Schneider, J., Lewen, A., Ta, T.T., Papageorgiou, I.E., and Kann, O. (2014). Energy substrates that fuel fast neuronal network oscillations. Front Neurosci. 8, 398.

German, M.S. (1993). Glucose sensing in pancreatic islet beta cells: the key role of glucokinase and the glycolytic intermediates. Proc. Natl. Acad. Sci. U. S. A 90, 1781–1785.

Gimenez-Cassina, A., Martinez-Francois, J.R., Fisher, J.K., Szlyk, B., Polak, K., Wiwczar, J., Tanner, G.R., Lutas, A., Yellen, G., and Danial, N.N. (2012). BAD-Dependent Regulation of Fuel Metabolism and K(ATP) Channel Activity Confers Resistance to Epileptic Seizures. Neuron 74, 719–730.

Girouard, H., Bonev, A.D., Hannah, R.M., Meredith, A., Aldrich, R.W., and Nelson, M.T. (2010). Astrocytic endfoot Ca2+ and BK channels determine both arteriolar dilation and constriction. Proc. Natl. Acad. Sci. U. S. A 107, 3811–3816.

Gribble, F.M., Ashfield, R., Ammala, C., and Ashcroft, F.M. (1997). Properties of cloned ATP-sensitive K+ currents expressed in Xenopus oocytes. J. Physiol 498 (Pt 1), 87–98.

Gulyas, A.I., Buzsaki, G., Freund, T.F., and Hirase, H. (2006). Populations of hippocampal inhibitory neurons express different levels of cytochrome c. Eur. J. Neurosci. 23, 2581–2594.

Gupta, A., Wang, Y., and Markram, H. (2000). Organizing principles for a diversity of GABAergic interneurons and synapses in the neocortex. Science 287, 273–278.

Haj-Dahmane, S., and Andrade, R. (1997). Calcium-activated cation nonselective current contributes to the fast afterdepolarization in rat prefrontal cortex neurons. J. Neurophysiol. 78, 1983–1989.

Halabisky, B.E., Shen, F., Huguenard, J.R., and Prince, D.A. (2006). Electrophysiological Classification of Somatostatin-positive Interneurons in Mouse Sensorimotor Cortex. J. Neurophysiol.

Hall, C.N., Klein-Flugge, M.C., Howarth, C., and Attwell, D. (2012). Oxidative phosphorylation, not glycolysis, powers presynaptic and postsynaptic mechanisms underlying brain information processing. J. Neurosci. 32, 8940–8951.

Heron-Milhavet, L., Xue-Jun, Y., Vannucci, S.J., Wood, T.L., Willing, L.B., Stannard, B., Hernandez-Sanchez, C., Mobbs, C., Virsolvy, A., and LeRoith, D. (2004). Protection against hypoxic-ischemic injury in transgenic mice overexpressing Kir6.2 channel pore in forebrain. Mol. Cell Neurosci. 25, 585–593.

Hill, E.L., Gallopin, T., Férézou, I., Cauli, B., Rossier, J., Schweitzer, P., and Lambolez, B. (2007). Functional CB1 receptors are broadly expressed in neocortical GABAergic and glutamatergic neurons. J. Neurophysiol. 97, 2580–2589.

Houades, V., Koulakoff, A., Ezan, P., Seif, I., and Giaume, C. (2008). Gap junction-mediated astrocytic networks in the mouse barrel cortex. J. Neurosci. 28, 5207–5217.

Hu, Y., and Wilson, G.S. (1997a). A temporary local energy pool coupled to neuronal activity: fluctuations of extracellular lactate levels in rat brain monitored with rapid-response enzyme-based sensor. J. Neurochem. 69, 1484–1490.

Hu, Y., and Wilson, G.S. (1997b). Rapid changes in local extracellular rat brain glucose observed with an in vivo glucose sensor. J. Neurochem. 68, 1745–1752.

Imamura, H., Nhat, K.P., Togawa, H., Saito, K., Iino, R., Kato-Yamada, Y., Nagai, T., and Noji, H. (2009). Visualization of ATP levels inside single living cells with fluorescence resonance energy transfer-based genetically encoded indicators. Proc. Natl. Acad. Sci. U. S. A 106, 15651–15656.

Inagaki, N., Gonoi, T., Clement, J.P., Namba, N., Inazawa, J., Gonzalez, G., guilar-Bryan, L., Seino, S., and Bryan, J. (1995a). Reconstitution of IKATP: an inward rectifier subunit plus the sulfonylurea receptor. Science 270, 1166–1170.

Inagaki, N., Gonoi, T., Clement, J.P., Wang, C.Z., Aguilar-Bryan, L., Bryan, J., and Seino, S. (1996). A family of sulfonylurea receptors determines the pharmacological properties of ATP-sensitive K+ channels. Neuron 16, 1011–1017.

Inagaki, N., Tsuura, Y., Namba, N., Masuda, K., Gonoi, T., Horie, M., Seino, Y., Mizuta, M., and Seino, S. (1995b). Cloning and functional characterization of a novel ATP-sensitive potassium channel ubiquitously expressed in rat tissues, including pancreatic islets, pituitary, skeletal muscle, and heart. J. Biol. Chem. 270, 5691–5694.

Isomoto, S., Kondo, C., Yamada, M., Matsumoto, S., Higashiguchi, O., Horio, Y., Matsuzawa, Y., and Kurachi, Y. (1996). A novel sulfonylurea receptor forms with BIR (Kir6.2) a smooth muscle type ATP-sensitive K+ channel. J. Biol. Chem. 271, 24321–24324.

Isomoto, S., and Kurachi, Y. (1997). Function, regulation, pharmacology, and molecular structure of ATP-sensitive K+ channels in the cardiovascular system. J. Cardiovasc. Electrophysiol. 8, 1431–1446.

Ivanov, A.I., Malkov, A.E., Waseem, T., Mukhtarov, M., Buldakova, S., Gubkina, O., Zilberter, M., and Zilberter, Y. (2014). Glycolysis and oxidative phosphorylation in neurons and astrocytes during network activity in hippocampal slices. J. Cereb. Blood Flow Metab 34, 397–407.

Jimenez-Blasco, D., Busquets-Garcia, A., Hebert-Chatelain, E., Serrat, R., Vicente-Gutierrez, C., Ioannidou, C., Gomez-Sotres, P., Lopez-Fabuel, I., Resch-Beusher, M., Resel, E., Arnouil, D., Saraswat, D., Varilh, M., Cannich, A., Julio-Kalajzic, F., Bonilla-Del, R., I, Almeida, A., Puente, N., Achicallende, S., Lopez-Rodriguez, M.L., Jolle, C., Deglon, N., Pellerin, L., Josephine, C., Bonvento, G., Panatier, A., Lutz, B., Piazza, P.V., Guzman, M., Bellocchio, L., Bouzier-Sore, A.K., Grandes, P., Bolanos, J.P., and Marsicano, G. (2020). Glucose metabolism links astroglial mitochondria to cannabinoid effects. Nature.

Kann, O., Papageorgiou, I.E., and Draguhn, A. (2014). Highly energized inhibitory interneurons are a central element for information processing in cortical networks. J. Cereb. Blood Flow Metab 34, 1270–1282.

Karagiannis, A., Gallopin, T., David, C., Battaglia, D., Geoffroy, H., Rossier, J., Hillman, E.M., Staiger, J.F., and Cauli, B. (2009). Classification of NPY-expressing neocortical interneurons. J. Neurosci. 29, 3642–3659.

Karagiannis, A., Sylantyev, S., Hadjihambi, A., Hosford, P.S., Kasparov, S., and Gourine, A.V. (2015). Hemichannel-mediated release of lactate. J. Cereb. Blood Flow Metab.

Karschin, C., Ecke, C., Ashcroft, F.M., and Karschin, A. (1997). Overlapping distribution of K(ATP) channel-forming Kir6.2 subunit and the sulfonylurea receptor SUR1 in rodent brain. FEBS Lett. 401, 59–64.

Kawaguchi, Y. (1993). Groupings of nonpyramidal and pyramidal cells with specific physiological and morphological characteristics in rat frontal cortex. J. Neurophysiol. 69, 416–431.

Kawaguchi, Y. (1995). Physiological subgroups of nonpyramidal cells with specific morphological characteristics in layer II/III of rat frontal cortex. J. Neurosci. 15, 2638–2655.

Kawamura, M., Jr., Ruskin, D.N., and Masino, S.A. (2010). Metabolic Autocrine Regulation of Neurons Involves Cooperation among Pannexin Hemichannels, Adenosine Receptors, and KATP Channels. J. Neurosci. 30, 3886–3895.

Krawchuk, M.B., Ruff, C.F., Yang, X., Ross, S.E., and Vazquez, A.L. (2019). Optogenetic assessment of VIP, PV, SOM and NOS inhibitory neuron activity and cerebral blood flow regulation in mouse somato-sensory cortex 1. J. Cereb. Blood Flow Metab 271678X19870105.

Lacroix, A., Toussay, X., Anenberg, E., Lecrux, C., Ferreiros, N., Karagiannis, A., Plaisier, F., Chausson, P., Jarlier, F., Burgess, S.A., Hillman, E.M., Tegeder, I., Murphy, T.H., Hamel, E., and Cauli, B. (2015). COX-2-Derived Prostaglandin E2 Produced by Pyramidal Neurons Contributes to Neurovascular Coupling in the Rodent Cerebral Cortex. J. Neurosci. 35, 11791–11810.

Lambolez, B., Audinat, E., Bochet, P., Crepel, F., and Rossier, J. (1992). AMPA receptor subunits expressed by single Purkinje cells. Neuron 9, 247–258.

Laughton, J.D., Charnay, Y., Belloir, B., Pellerin, L., Magistretti, P.J., and Bouras, C. (2000). Differential messenger RNA distribution of lactate dehydrogenase LDH-1 and LDH-5 isoforms in the rat brain. Neuroscience 96, 619–625.

Lauritzen, K.H., Morland, C., Puchades, M., Holm-Hansen, S., Hagelin, E.M., Lauritzen, F., Attramadal, H., Storm-Mathisen, J., Gjedde, A., and Bergersen, L.H. (2014). Lactate receptor sites link neurotransmission, neurovascular coupling, and brain energy metabolism. Cereb. Cortex 24, 2784–2795.

Le Douce J., Maugard, M., Veran, J., Matos, M., Jego, P., Vigneron, P.A., Faivre, E., Toussay, X., Vandenberghe, M., Balbastre, Y., Piquet, J., Guiot, E., Tran, N.T., Taverna, M., Marinesco, S., Koyanagi, A., Furuya, S., Gaudin-Guerif, M., Goutal, S., Ghettas, A., Pruvost, A., Bemelmans, A.P., Gaillard, M.C., Cambon, K., Stimmer, L., Sazdovitch, V., Duyckaerts, C., Knott, G., Herard, A.S., Delzescaux, T., Hantraye, P., Brouillet, E., Cauli, B., Oliet, S.H.R., Panatier, A., and Bonvento, G. (2020). Impairment of Glycolysis-Derived l-Serine Production in Astrocytes Contributes to Cognitive Deficits in Alzheimer’s Disease. Cell Metab 31, 503–517.

Lee, K.P.K., Chen, J., and MacKinnon, R. (2017). Molecular structure of human KATP in complex with ATP and ADP. Elife. 6.

Lee, Y., Morrison, B.M., Li, Y., Lengacher, S., Farah, M.H., Hoffman, P.N., Liu, Y., Tsingalia, A., Jin, L., Zhang, P.W., Pellerin, L., Magistretti, P.J., and Rothstein, J.D. (2012). Oligodendroglia metabolically support axons and contribute to neurodegeneration. Nature.

Lemak, M.S., Voloshanenko, O., Draguhn, A., and Egorov, A.V. (2014). KATP channels modulate intrinsic firing activity of immature entorhinal cortex layer III neurons. Front Cell Neurosci. 8, 255.

Lennie, P. (2003). The cost of cortical computation. Curr. Biol. 13, 493–497.

Lerchundi, R., Fernandez-Moncada, I., Contreras-Baeza, Y., Sotelo-Hitschfeld, T., Machler, P., Wyss, M.T., Stobart, J., Baeza-Lehnert, F., Alegria, K., Weber, B., and Barros, L.F. (2015). NH4+ triggers the release of astrocytic lactate via mitochondrial pyruvate shunting. Proc. Natl. Acad. Sci. U. S. A.

Li, N., Wu, J.X., Ding, D., Cheng, J., Gao, N., and Chen, L. (2017). Structure of a Pancreatic ATP-Sensitive Potassium Channel. Cell 168, 101–110.

Liss, B., Bruns, R., and Roeper, J. (1999). Alternative sulfonylurea receptor expression defines metabolic sensitivity of K-ATP channels in dopaminergic midbrain neurons. EMBO J. 18, 833–846.

Logothetis, N.K. (2008). What we can do and what we cannot do with fMRI. Nature 453, 869–878.

Lundgaard, I., Li, B., Xie, L., Kang, H., Sanggaard, S., Haswell, J.D., Sun, W., Goldman, S., Blekot, S., Nielsen, M., Takano, T., Deane, R., and Nedergaard, M. (2015). Direct neuronal glucose uptake heralds activity-dependent increases in cerebral metabolism. Nat. Commun. 6, 6807.

Machler, P., Wyss, M.T., Elsayed, M., Stobart, J., Gutierrez, R., von Faber-Castell, A., Kaelin, V., Zuend, M., San, M.A., Romero-Gomez, I., Baeza-Lehnert, F., Lengacher, S., Schneider, B.L., Aebischer, P., Magistretti, P.J., Barros, L.F., and Weber, B. (2016). In Vivo Evidence for a Lactate Gradient from Astrocytes to Neurons. Cell Metab 23, 94–102.

Magistretti, P.J., and Allaman, I. (2018). Lactate in the brain: from metabolic end-product to signalling molecule. Nat. Rev. Neurosci. 19, 235–249.

Martin, G.M., Yoshioka, C., Rex, E.A., Fay, J.F., Xie, Q., Whorton, M.R., Chen, J.Z., and Shyng, S.L. (2017). Cryo-EM structure of the ATP-sensitive potassium channel illuminates mechanisms of assembly and gating. Elife. 6.

Miki, T., Liss, B., Minami, K., Shiuchi, T., Saraya, A., Kashima, Y., Horiuchi, M., Ashcroft, F., Minokoshi, Y., Roeper, J., and Seino, S. (2001). ATP-sensitive K+ channels in the hypothalamus are essential for the maintenance of glucose homeostasis. Nat. Neurosci. 4, 507–512.

Miki, T., Nagashima, K., Tashiro, F., Kotake, K., Yoshitomi, H., Tamamoto, A., Gonoi, T., Iwanaga, T., Miyazaki, J., and Seino, S. (1998). Defective insulin secretion and enhanced insulin action in KATP channel-deficient mice. Proc. Natl. Acad. Sci. U. S. A 95, 10402–10406.

Molnar, G., Farago, N., Kocsis, A.K., Rozsa, M., Lovas, S., Boldog, E., Baldi, R., Csajbok, E., Gardi, J., Puskas, L.G., and Tamas, G. (2014). GABAergic neurogliaform cells represent local sources of insulin in the cerebral cortex. J. Neurosci. 34, 1133–1137.

Moreau, C., Prost, A.L., Derand, R., and Vivaudou, M. (2005). SUR, ABC proteins targeted by KATP channel openers. J. Mol. Cell Cardiol. 38, 951–963.

Newgard, C.B., and McGarry, J.D. (1995). Metabolic coupling factors in pancreatic beta-cell signal transduction. Annu. Rev. Biochem. 64, 689–719.

Ogawa, M., Watabe, H., Teramoto, N., Miyake, Y., Hayashi, T., Iida, H., Murata, T., and Magata, Y. (2005). Understanding of cerebral energy metabolism by dynamic living brain slice imaging system with [18F]FDG. Neurosci. Res. 52, 357–361.

Okuyama, Y., Yamada, M., Kondo, C., Satoh, E., Isomoto, S., Shindo, T., Horio, Y., Kitakaze, M., Hori, M., and Kurachi, Y. (1998). The effects of nucleotides and potassium channel openers on the SUR2A/Kir6.2 complex K+ channel expressed in a mammalian cell line, HEK293T cells. Pflugers Arch. 435, 595–603.

Pellerin, L., and Magistretti, P.J. (1994). Glutamate uptake into astrocytes stimulates aerobic glycolysis: a mechanism coupling neuronal activity to glucose utilization. Proc. Natl. Acad. Sci. U. S. A. 91, 10625–9.

Perrenoud, Q., Geoffroy, H., Gauthier, B., Rancillac, A., Alfonsi, F., Kessaris, N., Rossier, J., Vitalis, T., and Gallopin, T. (2012). Characterization of Type I and Type II nNOS-Expressing Interneurons in the Barrel Cortex of Mouse. Front Neural Circuits. 6, 36.

Piquet, J., Toussay, X., Hepp, R., Lerchundi, R., Le, D.J., Faivre, E., Guiot, E., Bonvento, G., and Cauli, B. (2018). Supragranular Pyramidal Cells Exhibit Early Metabolic Alterations in the 3xTg-AD Mouse Model of Alzheimer’s Disease. Front Cell Neurosci. 12, 216.

Prichard, J., Rothman, D., Novotny, E., Petroff, O., Kuwabara, T., Avison, M., Howseman, A., Hanstock, C., and Shulman, R. (1991). Lactate rise detected by 1H NMR in human visual cortex during physiologic stimulation. Proc. Natl. Acad. Sci. U. S. A 88, 5829–5831.

Puljung, M.C. (2018). Cryo-electron microscopy structures and progress toward a dynamic understanding of KATP channels. J. Gen. Physiol 150, 653–669.

Pullen, T.J., da, S., X, Kelsey, G., and Rutter, G.A. (2011). miR-29a and miR-29b contribute to pancreatic beta-cell-specific silencing of monocarboxylate transporter 1 (Mct1). Mol. Cell Biol. 31, 3182–3194.

Quistorff, B., Secher, N.H., and Van Lieshout, J.J. (2008). Lactate fuels the human brain during exercise. FASEB J. 22, 3443–3449.

Raichle, M.E., and Mintun, M.A. (2006). Brain work and brain imaging. Annu. Rev. Neurosci. 29, 449–476.

Rouach, N., Koulakoff, A., Abudara, V., Willecke, K., and Giaume, C. (2008). Astroglial metabolic networks sustain hippocampal synaptic transmission. Science 322, 1551–1555.

Ruminot, I., Gutierrez, R., Pena-Munzenmayer, G., Anazco, C., Sotelo-Hitschfeld, T., Lerchundi, R., Niemeyer, M.I., Shull, G.E., and Barros, L.F. (2011). NBCe1 Mediates the Acute Stimulation of Astrocytic Glycolysis by Extracellular K+. J. Neurosci. 31, 14264–14271.

Sada, N., Lee, S., Katsu, T., Otsuki, T., and Inoue, T. (2015). Epilepsy treatment. Targeting LDH enzymes with a stiripentol analog to treat epilepsy. Science 347, 1362–1367.

Sakura, H., Ammala, C., Smith, P.A., Gribble, F.M., and Ashcroft, F.M. (1995). Cloning and functional expression of the cDNA encoding a novel ATP-sensitive potassium channel subunit expressed in pancreatic beta-cells, brain, heart and skeletal muscle. FEBS Lett. 377, 338–344.

Saunders, A., Macosko, E.Z., Wysoker, A., Goldman, M., Krienen, F.M., de, R.H., Bien, E., Baum, M., Bortolin, L., Wang, S., Goeva, A., Nemesh, J., Kamitaki, N., Brumbaugh, S., Kulp, D., and McCarroll, S.A. (2018). Molecular Diversity and Specializations among the Cells of the Adult Mouse Brain. Cell 174, 1015–1030.

Schindelin, J., rganda-Carreras, I., Frise, E., Kaynig, V., Longair, M., Pietzsch, T., Preibisch, S., Rueden, C., Saalfeld, S., Schmid, B., Tinevez, J.Y., White, D.J., Hartenstein, V., Eliceiri, K., Tomancak, P., and Cardona, A. (2012). Fiji: an open-source platform for biological-image analysis. Nat. Methods 9, 676–682.

Schurr, A., Miller, J.J., Payne, R.S., and Rigor, B.M. (1999). An increase in lactate output by brain tissue serves to meet the energy needs of glutamate-activated neurons. J. Neurosci. 19, 34–39.

Schurr, A., West, C.A., and Rigor, B.M. (1988). Lactate-supported synaptic function in the rat hippocampal slice preparation. Science 240, 1326–1328.

Sekine, N., Cirulli, V., Regazzi, R., Brown, L.J., Gine, E., Tamarit-Rodriguez, J., Girotti, M., Marie, S., MacDonald, M.J., Wollheim, C.B., and. (1994). Low lactate dehydrogenase and high mitochondrial glycerol phosphate dehydrogenase in pancreatic beta-cells. Potential role in nutrient sensing. J. Biol. Chem. 269, 4895–4902.

Shmuel, A., Augath, M., Oeltermann, A., and Logothetis, N.K. (2006). Negative functional MRI response correlates with decreases in neuronal activity in monkey visual area V1. Nat. Neurosci. 9, 569–577.

Shmuel, A., Yacoub, E., Pfeuffer, J., Van de Moortele, P.F., Adriany, G., Hu, X., and Ugurbil, K. (2002). Sustained negative BOLD, blood flow and oxygen consumption response and its coupling to the positive response in the human brain. Neuron 36, 1195–1210.

Silver, I.A., and Erecinska, M. (1994). Extracellular glucose concentration in mammalian brain: continuous monitoring of changes during increased neuronal activity and upon limitation in oxygen supply in normo-, hypo-, and hyperglycemic animals. J. Neurosci. 14, 5068–5076.

Song, Z., and Routh, V.H. (2005). Differential effects of glucose and lactate on glucosensing neurons in the ventromedial hypothalamic nucleus. Diabetes 54, 15–22.

Sotelo-Hitschfeld, T., Niemeyer, M.I., Machler, P., Ruminot, I., Lerchundi, R., Wyss, M.T., Stobart, J., Fernandez-Moncada, I., Valdebenito, R., Garrido-Gerter, P., Contreras-Baeza, Y., Schneider, B.L., Aebischer, P., Lengacher, S., San, M.A., Le, D.J., Bonvento, G., Magistretti, P.J., Sepulveda, F.V., Weber, B., and Barros, L.F. (2015). Channel-mediated lactate release by k+-stimulated astrocytes. J. Neurosci. 35, 4168–4178.

Stella, N., Schweitzer, P., and Piomelli, D. (1997). A second endogenous cannabinoid that modulates long-term potentiation. Nature 388, 773–778.

Sun, H.S., Feng, Z.P., Miki, T., Seino, S., and French, R.J. (2006). Enhanced neuronal damage after ischemic insults in mice lacking Kir6.2-containing ATP-sensitive K+ channels. J. Neurophysiol. 95, 2590–2601.

Suzuki, A., Stern, S.A., Bozdagi, O., Huntley, G.W., Walker, R.H., Magistretti, P.J., and Alberini, C.M. (2011). Astrocyte-neuron lactate transport is required for long-term memory formation. Cell 144, 810–823.

Tanner, G.R., Lutas, A., Martinez-Francois, J.R., and Yellen, G. (2011). Single K ATP channel opening in response to action potential firing in mouse dentate granule neurons. J. Neurosci. 31, 8689–8696.

Tantama, M., Martinez-Francois, J.R., Mongeon, R., and Yellen, G. (2013). Imaging energy status in live cells with a fluorescent biosensor of the intracellular ATP-to-ADP ratio. Nat. Commun. 4, 2550.

Tarasov, A.I., Girard, C.A., and Ashcroft, F.M. (2006). ATP sensitivity of the ATP-sensitive K+ channel in intact and permeabilized pancreatic beta-cells. Diabetes 55, 2446–2454.

Tasic, B., Menon, V., Nguyen, T.N., Kim, T.K., Jarsky, T., Yao, Z., Levi, B., Gray, L.T., Sorensen, S.A., Dolbeare, T., Bertagnolli, D., Goldy, J., Shapovalova, N., Parry, S., Lee, C., Smith, K., Bernard, A., Madisen, L., Sunkin, S.M., Hawrylycz, M., Koch, C., and Zeng, H. (2016). Adult mouse cortical cell taxonomy revealed by single cell transcriptomics. Nat. Neurosci.

Thevenaz, P., Ruttimann, U.E., and Unser, M. (1998). A pyramid approach to subpixel registration based on intensity. IEEE Trans. Image Process 7, 27–41.

Thoby-Brisson, M., Cauli, B., Champagnat, J., Fortin, G., and Katz, D.M. (2003). Expression of functional tyrosine kinase B receptors by rhythmically active respiratory neurons in the pre-Botzinger complex of neonatal mice. J. Neurosci. 23, 7685–7689.

Thomzig, A., Laube, G., Pruss, H., and Veh, R.W. (2005). Pore-forming subunits of K-ATP channels, Kir6.1 and Kir6.2, display prominent differences in regional and cellular distribution in the rat brain. J. Comp Neurol. 484, 313–330.

Tsuzuki, K., Lambolez, B., Rossier, J., and Ozawa, S. (2001). Absolute quantification of AMPA receptor subunit mRNAs in single hippocampal neurons. J. Neurochem. 77, 1650–1659.

Uhlirova, H., Kilic, K., Tian, P., Thunemann, M., Desjardins, M., Saisan, P.A., Sakadzic, S., Ness, T.V., Mateo, C., Cheng, Q., Weldy, K.L., Razoux, F., Vanderberghe, M., Cremonesi, J.A., Ferri, C.G., Nizar, K., Sridhar, V.B., Steed, T.C., Abashin, M., Fainman, Y., Masliah, E., Djurovic, S., Andreassen, O., Silva, G.A., Boas, D.A., Kleinfeld, D., Buxton, R.B., Einevoll, G.T., Dale, A.M., and Devor, A. (2016). Cell type specificity of neurovascular coupling in cerebral cortex. Elife. 5.

Vanlandewijck, M., He, L., Mae, M.A., Andrae, J., Ando, K., Del, G.F., Nahar, K., Lebouvier, T., Lavina, B., Gouveia, L., Sun, Y., Raschperger, E., Rasanen, M., Zarb, Y., Mochizuki, N., Keller, A., Lendahl, U., and Betsholtz, C. (2018). A molecular atlas of cell types and zonation in the brain vasculature. Nature 554, 475–480.

Varin, C., Rancillac, A., Geoffroy, H., Arthaud, S., Fort, P., and Gallopin, T. (2015). Glucose Induces Slow-Wave Sleep by Exciting the Sleep-Promoting Neurons in the Ventrolateral Preoptic Nucleus: A New Link between Sleep and Metabolism. J. Neurosci. 35, 9900–9911.

Vezzoli, E., Cali, C., De, R.M., Ponzoni, L., Sogne, E., Gagnon, N., Francolini, M., Braida, D., Sala, M., Muller, D., Falqui, A., and Magistretti, P.J. (2020). Ultrastructural Evidence for a Role of Astrocytes and Glycogen-Derived Lactate in Learning-Dependent Synaptic Stabilization. Cereb. Cortex 30, 2114–2127.

Voutsinos-Porche, B., Bonvento, G., Tanaka, K., Steiner, P., Welker, E., Chatton, J.Y., Magistretti, P.J., and Pellerin, L. (2003). Glial glutamate transporters mediate a functional metabolic crosstalk between neurons and astrocytes in the mouse developing cortex. Neuron 37, 275–286.

Ward, J.H. (1963). Hierarchical grouping to optimize an objective function. Journal of the American Statistical Association 58, 236–244.

Wilson, J.E. (2003). Isozymes of mammalian hexokinase: structure, subcellular localization and metabolic function. J. Exp. Biol. 206, 2049–2057.

Wyss, M.T., Jolivet, R., Buck, A., Magistretti, P.J., and Weber, B. (2011). In vivo evidence for lactate as a neuronal energy source. J. Neurosci. 31, 7477–7485.

Xi, Q., Cheranov, S.Y., and Jaggar, J.H. (2005). Mitochondria-derived reactive oxygen species dilate cerebral arteries by activating Ca2+ sparks. Circ. Res. 97, 354–362.

Yamada, M., Isomoto, S., Matsumoto, S., Kondo, C., Shindo, T., Horio, Y., and Kurachi, Y. (1997). Sulphonylurea receptor 2B and Kir6.1 form a sulphonylurea-sensitive but ATP-insensitive K+ channel. J. Physiol 499 (Pt 3), 715–720.

Yang, X.J., Kow, L.M., Funabashi, T., and Mobbs, C.V. (1999). Hypothalamic glucose sensor: similarities to and differences from pancreatic beta-cell mechanisms. Diabetes 48, 1763–1772.

Zawar, C., and Neumcke, B. (2000). Differential activation of ATP-sensitive potassium channels during energy depletion in CA1 pyramidal cells and interneurones of rat hippocampus. Pflugers Arch. 439, 256–262.

Zawar, C., Plant, T.D., Schirra, C., Konnerth, A., and Neumcke, B. (1999). Cell-type specific expression of ATP-sensitive potassium channels in the rat hippocampus. J. Physiol 514 (Pt 2), 327–341.

Zeisel, A., Manchado, A.B., Codeluppi, S., Lonnerberg, P., La, M.G., Jureus, A., Marques, S., Munguba, H., He, L., Betsholtz, C., Rolny, C., Castelo-Branco, G., Hjerling-Leffler, J., and Linnarsson, S. (2015). Cell types in the mouse cortex and hippocampus revealed by single-cell RNA-seq. Science.

