## Supplementary tables S1-7 for "Lactate is a major energy substrate for cortical neurons and enhances their firing activity"

### Table S1. Somatic properties of different neuronal types

| | **RS** | **IB** | **Burst. VIP** | **Adapt. VIP** | **Adapt. SOM** | **Adapt. NPY** | **FS-PV** |
|---|---|---|---|---|---|---|---|
| | **n = 63** | **n = 10** | **n = 27** | **n = 59** | **n = 24** | **n = 56** | **n = 38** |
| **Layer** | 2.9 ± 0.1 | **3.9 ± 0.1** | 2.5 ± 0.1 | 2.6 ± 0.1 | 3.0 ± 0.1 | **2.1 ± 0.1** | 3.1 ± 0.1 |
| | **Adapt. NPY** <<< RS, IB, Adapt. VIP, Adapt. SOM, FS-PV<br>Burst. VIP, Adapt. VIP <<< **IB**; Adapt. SOM << **IB**; RS < **IB** | | | | | | |
| **Major axis (µm)** | 22.3 ± 1.2 | 16.7 ± 1.3 | 17.8 ± 0.9 | 20.5 ± 0.6 | 24.7 ± 1.6 | 21.2 ± 1.1 | 22.9 ± 1.1 |
| | IB < Adapt. SOM, FS-PV | | | | | | |
| **Minor axis (µm)** | 11.5 ± 0.5 | 9.2 ± 0.3 | 9.2 ± 0.3 | 9.0 ± 0.2 | 9.4 ± 0.3 | 9.6 ± 0.2 | 9.8 ± 0.3 |
| | Adapt. VIP <<< RS | | | | | | |
| **Elongation** | 2.0 ± 0.1 | 1.8 ± 0.2 | 2.0 ± 0.1 | 2.3 ± 0.1 | 2.6 ± 0.2 | 2.2 ± 0.1 | 2.3 ± 0.1 |
| | RS << Adapt. VIP, Adapt. SOM; RS < FS-PV<br>IB < Adapt. VIP, Adapt. SOM, FS-PV | | | | | | |
| **Area (µm$^2$)** | 157.3 ± 8.3 | 115.4 ± 6.9 | 116.0 ± 6.4 | 127.0 ± 4.2 | 161.6 ± 10.9 | 145.0 ± 7.6 | 160.9 ± 7.0 |
| | Burst. VIP, Adapt. VIP <<< FS-PV; IB < FS-PV<br>Burst. VIP << Adapt. SOM; IB, Adapt. VIP < Adapt. SOM<br>Burst. VIP << RS | | | | | | |
| **Perimeter (µm)** | 58.0 ± 2.9 | 44.2 ± 2.3 | 44.7 ± 1.8 | 48.8 ± 1.2 | 58.0 ± 3.3 | 52.1 ± 2.1 | 54.9 ± 2.5 |
| | IB, Burst. VIP, Adapt. VIP < FS-PV | | | | | | |
| **Roundness** | 1.7 ± 0.1 | 1.4 ± 0.1 | 1.4 ± 0.0 | 1.6 ± 0.1 | 1.7 ± 0.1 | 1.5 ± 0.0 | 1.6 ± 0.1 |
| | n.s. | | | | | | |

n, number of cells, < significantly smaller with $P \leq 0.05$; << significantly smaller with $P \leq 0.01$; <<< significantly smaller with $P \leq 0.001$. . n.s. not statistically significant.

### Table S2. Occurrence of molecular markers in different neuronal types

<table>
<tr><th></th><th>RS<br>n = 63</th><th>IB<br>n = 10</th><th>Burst. VIP<br>n = 27</th><th>Adapt. VIP<br>n = 59</th><th>Adapt. SOM<br>n = 24</th><th>Adapt. NPY<br>n = 56</th><th>FS-PV<br>n = 38</th></tr>
<tr><td rowspan="2">VGluT1</td><td>94</td><td>100</td><td>44</td><td>17</td><td>33</td><td>34</td><td>32</td></tr>
<tr><td colspan="7">RS >>>Burst. VIP, Adapt. VIP, Adapt. SOM, Adapt. NPY, FS,PV<br>IB >>> Adapt. VIP; IB >> Adapt. SOM, Adapt. NPY, FS-PV; IB > Burst. VIP</td></tr>
<tr><td rowspan="2">GAD</td><td>10</td><td>10</td><td>100</td><td>100</td><td>100</td><td>96</td><td>100</td></tr>
<tr><td colspan="7">Burst. VIP, Adapt. VIP, Adapt. SOM, Adapt. NPY, FS,PV >>> RS, IB</td></tr>
<tr><td rowspan="2">NOS-1</td><td>0</td><td>0</td><td>0</td><td>5</td><td>13</td><td>21</td><td>8</td></tr>
<tr><td colspan="7">Adapt. NPY >> RS</td></tr>
<tr><td rowspan="2">CB</td><td>40</td><td>40</td><td>7</td><td>17</td><td>88</td><td>11</td><td>50</td></tr>
<tr><td colspan="7">Adapt. SOM >>> Burst. VIP, Adapt. VIP, Adapt. NPY; Adapt. SOM >> RS, Adapt. SOM > FS-PV<br>FS-PV >>> Adapt. NPY; FS-PV >> Burst. VIP, Adapt. VIP<br>RS >> Adapt. NPY, RS > Burst. VIP</td></tr>
<tr><td rowspan="2">PV</td><td>33</td><td>20</td><td>11</td><td>20</td><td>38</td><td>21</td><td>97</td></tr>
<tr><td colspan="7">FS-PV >>> RS, IB, Burst. VIP, Adapt. VIP, Adapt. NPY; Adapt. SOM</td></tr>
<tr><td rowspan="2">CR</td><td>0</td><td>0</td><td>26</td><td>39</td><td>4</td><td>16</td><td>3</td></tr>
<tr><td colspan="7">Adapt. VIP >>> RS, FS-PV, Adapt. VIP > Adapt. SOM<br>Burst. VIP >> RS, Adapt. NPY > RS</td></tr>
<tr><td rowspan="2">NPY</td><td>0</td><td>10</td><td>7</td><td>12</td><td>67</td><td>55</td><td>24</td></tr>
<tr><td colspan="7">Adapt. SOM, Adapt. NPY >>> RS, Burst. VIP, Adapt. VIP; Adapt. SOM, Adapt. NPY > FS-PV<br>FS-PV >> RS</td></tr>
<tr><td rowspan="2">VIP</td><td>0</td><td>0</td><td>89</td><td>83</td><td>4</td><td>11</td><td>3</td></tr>
<tr><td colspan="7">Burst. VIP, Adapt. VIP >>> RS, IB, Adapt. SOM, Adapt. NPY, FS-PV</td></tr>
<tr><td rowspan="2">SOM</td><td>10</td><td>0</td><td>11</td><td>8</td><td>92</td><td>9</td><td>5</td></tr>
<tr><td colspan="7">Adapt. SOM >>> RS, IB, Burst. VIP, Adapt. VIP, Adapt. NPY, FS-PV</td></tr>
<tr><td rowspan="2">CCK</td><td>33</td><td>10</td><td>22</td><td>34</td><td>4</td><td>18</td><td>8</td></tr>
<tr><td colspan="7">n.s.</td></tr>
</table>

Occurrences are given in %; n, number of cells; > significantly larger with $P \leq 0.05$; >> significantly larger with $P \leq 0.01$; >>> significantly larger with $P \leq 0.001$. n.s. not statistically significant.

### Table S3. Passive properties of different neuronal types

| | RS | IB | Burst. VIP | Adapt. VIP | Adapt. SOM | Adapt. NPY | FS-PV |
|---|---|---|---|---|---|---|---|
| | **n = 63** | **n = 10** | **n = 27** | **n = 59** | **n = 24** | **n = 56** | **n = 38** |
| **(1) Resting potential (mV)** | **-81.5 ± 0.9** | -79.2 ± 1.1 | -75.6 ± 1.2 | -73.9 ± 0.8 | **-71.6 ± 1.1** | -76.6 ± 0.7 | -77.4 ± 0.9 |
| | RS <<< Adapt. VIP, Adapt. SOM; RS << Adapt. NPY; RS < Burst. VIP, FS-PV<br>IB, Adapt. NPY << Adapt. SOM; FS-PV <<< Adapt. SOM | | | | | | |
| **(2) Input resistance (MΩ)** | 319 ± 7 | 361 ± 39 | **608 ± 64** | **569 ± 26** | 350 ± 36 | 387 ± 22 | **201 ± 13** |
| | **FS-PV** <<< RS, IB, Burst. VIP, Adapt. VIP, Adapt. SOM, Adapt. NPY<br>RS, Adapt. SOM, Adapt. NPY <<< **Adapt. VIP**; IB << **Adapt. VIP**<br>RS <<< **Burst. VIP;** Adapt. SOM, Adapt. NPY << **Burst. VIP** | | | | | | |
| **(3) Time constant (ms)** | 32.4 ± 1.4 | 33.5 ± 3.1 | 31.0 ± 2.6 | 27.8 ± 1.5 | 28.3 ± 3.3 | 28.1 ± 1.5 | **16.2 ± 1.0** |
| | **FS-PV** <<< RS, IB, Burst. VIP, Adpat. VIP, Adapt. NPY; **FS-PV** << Adapt. SOM | | | | | | |
| **(4) Membrane capacitance (pF)** | **110.7 ± 4.1** | 95.6 ± 5.9 | **56.3 ± 4.0** | **51.1 ± 2.8** | 82.8 ± 4.7 | 76.8 ± 3.3 | 83.8 ± 3.6 |
| | **Adapt. VIP** <<< RS, IB, Adapt. SOM, Adapt. NPY, FS-PV<br>**Burst. VIP** <<< RS, IB, Adapt. SOM, FS-PV; **Burst. VIP** << Adapt. NPY<br>Adapt. SOM, Adapt. NPY, FS-PV <<< **RS** | | | | | | |
| **(5) Sag index (%)** | 14.1 ± 1.1 | **26.1 ± 2.3** | 5.7 ± 0.9 | 8.3 ± 0.7 | **24.9 ± 2.5** | 9.5 ± 0.6 | 9.0 ± 0.9 |
| | Burst. VIP, Adapt. VIP, Adapt. NPY, FS-PV <<< **IB, Adapt. SOM**; RS << **IB**, **Adapt. SOM**<br>Burst. VIP <<< RS; Burst. VIP << Adapt. NPY<br>Adapt. VIP << RS; FS-PV < RS | | | | | | |

n, number of cells, < significantly smaller with P ≤ 0.05; << significantly smaller with P ≤ 0.01; <<< significantly smaller with P ≤ 0.001

### Table S4. Just above threshold properties of different neuronal types

| | RS | IB | Burst. VIP | Adapt. VIP | Adapt. SOM | Adapt. NPY | FS-PV |
|---|---|---|---|---|---|---|---|
| | **n = 63** | **n = 10** | **n = 27** | **n = 59** | **n = 24** | **n = 56** | **n = 38** |
| **(6) Rheobase (pA)** | 52.0 ± 4.2 | 39.2 ± 7.5 | 25.8 ± 4.1 | **14.5 ± 2.3** | **14.0 ± 6.2** | 40.8 ± 4.1 | **99.3 ± 8.4** |
| | RS, IB, Burst. VIP, Adapt VIP, Adapt. SOM, Adapt. NPY <<< **FS-PV**<br>**Adapt. VIP** <<< RS, Adapt. NPY; **Adapt. VIP** << IB<br>**Adapt. SOM** <<< RS; **Adapt. SOM** << Adapt. NPY<br>Burst. VIP << RS | | | | | | |
| **(7) First spike latency (ms)** | 151.3 ± 9.5 | 147.9 ± 25.2 | 102.2 ± 19.3 | 109.4 ± 10.7 | 144.3 ± 27.2 | 224.9 ± 26.1 | **348.3 ± 42.2** |
| | Burst. VIP, Adapt. VIP <<< **FS-PV**; RS, Adapt. SOM << **FS-PV**<br>Adapt. VIP <<< RS, Burst. VIP << RS<br>Adapt. VIP, Burst. VIP << Adapt. NPY | | | | | | |
| **(8) Adaptation (Hz/s)** | -9.8 ± 2.9 | **-168.8 ± 26.0** | **-658.4 ± 412.9** | -8.3 ± 3.0 | -14.4 ± 4.1 | **-1.9 ± 0.7** | **20.9 ± 22.2** |
| | **IB, Burst. VIP** <<< RS, Adapt. VIP, Adapt. SOM, Adapt. NPY, FS-PV<br>RS, Adapt. VIP, Adapt. SOM << **Adapt. NPY, FS-PV** | | | | | | |
| **(9) Minimal steady state frequency (Hz)** | 8.2 ± 1.3 | **66.2 ± 4.8** | **61.4 ± 8.4** | 11.6 ± 0.9 | 13.3 ± 2.3 | 6.8 ± 0.4 | 15.1 ± 1.1 |
| | RS, Adapt. VIP, Adapt. SOM, Adapt. NPY, FS-PV <<< **IB, Burst. VIP**<br>**Adapt. NPY** <<< Adapt. VIP, FS-PV, **Adapt. NPY** << Adapt. SOM<br>RS <<< Adapt. VIP << FS-PV | | | | | | |

n, number of cells; < significantly smaller with P ≤ 0.05; << significantly smaller with P ≤ 0.01; <<< significantly smaller with P ≤ 0.001

### Table S5. Firing properties of different neuronal types

| | RS | IB | Burst. VIP | Adapt. VIP | Adapt. SOM | Adapt. NPY | FS-PV |
|---|---|---|---|---|---|---|---|
| | n = 63 | n = 10 | n = 27 | n = 59 | n = 24 | n = 56 | n = 38 |
| **(10) Amplitude accommodation (mV)** | **18.5 ± 1.1** | **36.7 ± 3.1** | 9.9 ± 1.4 | 5.7 ± 0.5 | 3.8 ± 0.6 | 8.7 ± 0.9 | **2.0 ± 0.3** |
| | Burst. VIP, Adapt. VIP, Adapt. SOM, Adapt. NPY, FS-PV <<< **RS** <<< **IB**<br>**FS-PV** <<< Burst. VIP, Adapt. VIP, Adapt NPY; FS-PV < Adapt. SOM<br>Adapt. SOM << Burst. VIP, Adapt. NPY; Adapt. SOM < Adapt. VIP<br>Adapt. VIP < Burst. VIP, Adapt. NPY | | | | | | |
| **(11) Amplitude of early adaptation (Hz)** | 85.2 ± 3.9 | 96.1 ± 6.4 | 79.9 ± 8.3 | 76.1 ± 4.3 | 76.9 ± 7.4 | 84.6 ± 3.3 | **47.9 ± 3.1** |
| | **FS-PV** <<< RS, IB, Burst. VIP, Adapt. VIP, Adapt. NPY; **FS-PV** << Adapt. SOM | | | | | | |
| **(12) Time constant of early adaptation (ms)** | **19.6 ± 0.8** | **38.5 ± 5.2** | 25.8 ± 2.4 | 26.0 ± 1.3 | **38.6 ± 3.7** | 25.6 ± 1.0 | 21.1 ± 3.1 |
| | **RS** <<< IB, Adapt. SOM, Adapt NPY; **RS** << Adapt. VIP, **RS** < Burst. VIP<br>Adapt. NPY, FS-PV <<< **Adapt. SOM**; Adapt. VIP << **Adapt. SOM**; Burst. VIP < **Adapt. SOM**<br>Adapt. NPY << **IB**; Burst. VIP, Adapt. VIP, FS-PV < **IB**<br>FS-PV < Adapt. VIP, Adapt. NPY | | | | | | |
| **(13) Late adaptation (Hz/s)** | -16.2 ± 1.4 | **-3.6 ± 1.8** | -204.1 ± 181.7 | -35.7 ± 2.9 | -24.8 ± 2.9 | -19.5 ± 1.2 | -29.4 ± 2.1 |
| | **IB** << RS; **IB** <<< Burst. VIP, Adapt. VIP, Adapt. SOM, Adapt . NPY, FS-PV<br>RS, Adapt. NPY <<< Adapt. VIP, FS-PV | | | | | | |
| **(14) Maximal steady state frequency (Hz)** | **37.2 ± 1.7** | **24.4 ± 2.5** | 68.7 ± 7.3 | 80.0 ± 5.6 | 72.8 ± 5.2 | 56.4 ± 2.0 | **139.9 ± 6.8** |
| | **IB** << **RS** <<< Burst. VIP, Adapt. VIP, Adapt. SOM, Adapt. NPY <<< **FS-PV**<br>Adapt. NPY << Adapt. VIP; Adapt. NPY < Adapt. SOM | | | | | | |

n, number of cells; < significantly smaller with P ≤ 0.05; << significantly smaller with P ≤ 0.01; <<< significantly smaller with P ≤ 0.001

### Table S6. Action potentials properties of different neuronal types

| | RS | IB | Burst. VIP | Adapt. VIP | Adapt. SOM | Adapt. NPY | FS-PV |
|---|---|---|---|---|---|---|---|
| | n = 63 | n = 10 | n = 27 | n = 59 | n = 24 | n = 56 | n = 38 |
| **(15) First spike amplitude (mV)** | 92.1 ± 1.2 | 90.8 ± 1.5 | 94.3 ± 2.3 | 90.9 ± 1.3 | 94.9 ± 2.5 | 89.0 ± 1.3 | **80.9 ± 1.4** |
| | **FS-PV** <<< RS, Burst. VIP, Adapt. VIP, Adapt. SOM, Adapt. NPY; **FS-PV** << IB | | | | | | |
| **(16) First spike duration (ms)** | **1.4 ± 0.0** | 1.4 ± 0.1 | 0.9 ± 0.1 | 0.9 ± 0.0 | 0.9 ± 0.1 | 1.1 ± 0.0 | **0.6 ± 0.0** |
| | **FS-PV** <<< Burst. VIP, Adapt. VIP, Adapt. SOM, Adapt. NPY <<< **RS**<br>**FS-PV** <<< **IB**; Burst. VIP, Adapt. VIP, Adapt. SOM <<< **IB**, Adapt. NPY << **IB**<br>Burst. VIP, Adapt. VIP <<< Adapt. NPY; Adapt. SOM < Adapt. NPY | | | | | | |
| **(17) Second spike amplitude (mV)** | 89.6 ± 1.3 | **68.4 ± 1.8** | **75.5 ± 2.1** | 87.1 ± 1.4 | 91.9 ± 2.2 | 86.0 ± 1.3 | **79.9 ± 1.5** |
| | **IB, Burst. VIP** <<< RS, Adapt. VIP, Adapt. SOM, Adapt. NPY, **IB** << **FS-PV**<br>**FS-PV** <<< RS, Adapt. SOM; **FS-PV** << Adapt. VIP, Adapt. VIP | | | | | | |
| **(18) Second spike duration (ms)** | **1.5 ± 0.0** | **1.8 ± 0.1** | 1.0 ± 0.1 | 0.9 ± 0.0 | 1.0 ± 0.1 | 1.2 ± 0.0 | **0.6 ± 0.0** |
| | **FS-PV** <<< Burst. VIP, Adapt. VIP, Adapt. SOM, Adapt. NPY <<< **RS** << **IB**<br>Adapt. VIP <<< Adapt. NPY; Burst. VIP, Adapt. SOM < Adapt. NPY | | | | | | |
| **(19) Amplitude Reduction (%)** | 2.8 ± 0.4 | **24.7 ± 1.5** | **19.7 ± 1.7** | 4.2 ± 0.4 | 2.9 ± 0.7 | 3.3 ± 0.6 | **1.2 ± 0.9** |
| | RS, Adapt. VIP, Adapt. SOM, Adapt. NPY<<< **IB, Burst. VIP**<br>**FS-PV** <<< Adapt.VIP; **FS-PV** < RS, Adapt. NPY, RS < Adapt. VIP | | | | | | |
| **(20) Duration Increase (%)** | 4.8 ± 0.4 | **31.3 ± 3.4** | **14.5 ± 1.8** | 4.5 ± 0.5 | 4.9 ± 0.6 | 7.6 ± 0.9 | **0.8 ± 0.5** |
| | **FS-PV** <<< Adapt. VIP, Adapt. SOM, Adapt. NPY<<< **Burst .VIP** <<< **IB**<br>Adapt. VIP << Adapt. NPY, RS < Adapt. NPY | | | | | | |

n, number of cells; < significantly smaller with P ≤ 0.05; << significantly smaller with P ≤ 0.01; <<< significantly smaller with P ≤ 0.001

### Table S7. AH and AD properties of different neuronal types

| | RS | IB | Burst. VIP | Adapt. VIP | Adapt. SOM | Adapt. NPY | FS-PV |
|---|---|---|---|---|---|---|---|
| | **n = 63** | **n = 10** | **n = 27** | **n = 59** | **n = 24** | **n = 56** | **n = 38** |
| **(21) First spike, fast AH (mV)** | -7.2 ± 0.3 | -7.0 ± 0.8 | -12.3 ± 0.7 | -14.7 ± 0.6 | -14.2 ± 1.1 | -14.5 ± 0.7 | **-23.7 ± 0.6** |
| | **FS-PV** <<< Burst. VIP, Adapt. VIP, Adapt. SOM, Adapt. NPY <<< **RS, IB** | | | | | | |
| **(22) first spike AD (mV)** | 1.0 ± 0.2 | 0.0 ± 0.0 | 0.1 ± 0.1 | **5.2 ± 0.5** | **3.1 ± 0.6** | 0.3 ± 0.1 | 0.8 ± 0.3 |
| | RS, IB, Burst. VIP, Adapt. NPY, FS-PV <<< **Adapt. VIP**<br>Burst. VIP, Adapt. NPY <<< **Adapt. SOM,** RS, IB, FS-PV <<< **Adapt. SOM** | | | | | | |
| **(23) First spike, medium AP (mV)** | **-14.5 ± 0.4** | **0.0 ± 0.0** | **-0.3 ± 0.3** | -10.9 ± 0.6 | -8.2 ± 1.4 | -11.6 ± 1.0 | -4.5 ± 1.4 |
| | **RS** <<< IB, Burst. VIP, Adapt. VIP, Adapt. SOM, FS-PV<br>Adapt. VIP, Adapt. NPY <<< **IB, Burst. VIP**, Adapt. SOM <<< **Burst. VIP**, Adapt. SOM << **IB**<br>Adapt .VIP <<< FS-PV, Adapt. NPY << FS-PV | | | | | | |
| **(24) First spike, fast AP latency (ms)** | **6.5 ± 0.3** | 4.4 ± 0.2 | 2.8 ± 0.2 | 3.2 ± 0.2 | 3.3 ± 0.2 | 5.2 ± 0.2 | 2.7 ± 0.2 |
| | Burst. VIP, Adapt. VIP, Adapt. SOM, FS-PV <<< Adapt. NPY << **RS**<br>Burst. VIP, FS-PV <<< IB; Adapt. VIP << IB, IB < Adapt. SOM, **RS** | | | | | | |
| **(25) First spike, AD latency (ms)** | 4.6 ± 0.7 | 0.0 ± 0.0 | 0.4 ± 0.4 | **9.9 ± 0.7** | **7.5 ± 1.3** | 3.1 ± 0.6 | 1.2 ± 0.4 |
| | RS, IB, Burst. VIP, Adapt. NPY, FS-PV <<< **Adapt. VIP**<br>Burst. VIP, FS-PV <<< **Adapt. SOM**; IB, Adapt. NPY < **Adapt. SOM** | | | | | | |
| **(26) First spike, medium AH latency (ms)** | **63.1 ± 3.0** | 0.0 ± 0.0 | 0.5 ± 0.5 | **30.0 ± 2.5** | 23.1 ± 3.9 | 18.5 ± 1.9 | 2.2 ± 0.7 |
| | IB, Burst. VIP, Adapt. VIP, Adapt. SOM, Adapt. NPY, FS-PV <<< **RS**<br>IB, Burst. VIP, FS-PV <<< Adapt. NPY << **Adapt. VIP**<br>Burst. VIP, Adapt. NPY <<< Adapt. SOM, IB << Adapt. SOM | | | | | | |
| **(27) Second spike, fast AH (mV)** | **-7.7 ± 0.3** | **-8.7 ± 0.8** | -14.3 ± 0.7 | -15.9 ± 0.6 | -13.8 ± 1.1 | -15.3 ± 0.8 | **-24.0 ± 0.6** |
| | **FS-PV** <<< Burst. VIP, Adapt. VIP Adapt. SOM, Adapt. NPY <<< **RS**<br>**FS-PV**, Burst. VIP, Adapt. VIP Adapt. NPY <<< **IB**, Adapt. SOM< **IB** | | | | | | |
| **(28) Second spike AD (mV)** | 0.2 ± 0.0 | 0.4 ± 0.2 | 2.8 ± 0.7 | **4.3 ± 0.4** | 2.8 ± 0.5 | 0.1 ± 0.0 | 0.8 ± 0.3 |
| | RS, IB, Adapt. NPY, FS-PV <<< **Adapt. VIP**<br>RS, Adapt NPY <<< Adapt. SOM; FS-PV << Adapt. SOM | | | | | | |
| **(29) Second spike, medium AH (mV)** | **-17.3 ± 0.5** | **-24.9 ± 0.9** | -6.9 ± 1.4 | -12.7 ± 0.6 | -8.1 ± 1.3 | -13.2 ± 1.1 | -4.0 ± 1.3 |
| | **IB** <<< **RS** <<< Burst. VIP, Adapt. VIP Adapt. SOM, Adapt. NPY, FS-PV<br>Adapt. NPY, Adapt VIP << FS-PV; Adapt. NPY << Burst. VIP; Adapt VIP < Burst. VIP<br>Adapt. NPY, Adapt VIP < Adapt. SOM; Adapt. SOM < FS-PV | | | | | | |
| **(30) Second spike, fast AH latency (ms)** | **6.9 ± 0.3** | **7.5 ± 0.5** | 3.5 ± 0.3 | 3.6 ± 0.2 | 3.5 ± 0.3 | 5.6 ± 0.2 | **2.8 ± 0.2** |
| | **FS-PV** < Burst. VIP, Adapt. VIP, Adapt. SOM <<< Adapt. NPY << **RS, IB** | | | | | | |
| **(31) Second spike AD latency (ms)** | 1.8 ± 0.4 | 6.6 ± 1.6 | 4.3 ± 1.0 | **8.7 ± 0.6** | **7.5 ± 1.2** | 1.6 ± 0.4 | 0.9 ± 0.3 |
| | RS, Adapt. NPY, FS-PV<<< **Adapt. VIP, Adapt. SOM**; Burst. VIP < **Adapt. VIP**<br>RS, Adapt. NPY, FS-PV< IB | | | | | | |
| **(32) Second spike, medium AH latency (ms)** | **62.9 ± 2.9** | **100.0 ± 8.0** | 21.9 ± 4.8 | 28.8 ± 2.1 | 24.0 ± 3.8 | 19.4 ± 2.2 | **2.1 ± 0.7** |
| | **FS-PV** <<< Adapt.VIP, Adapt. SOM, Adapt. NPY <<< **RS** <<< **IB**<br>Burst. VIP <<< **RS, IB;** FS-PV < Burst. VIP; Adapt. NPY << Adapt. VIP | | | | | | |

n, number of cells; < significantly smaller with $P \leq 0.05$; << significantly smaller with $P \leq 0.01$; <<< significantly smaller with $P \leq 0.001$
